# ATG6 interacting with NPR1 increases *Arabidopsis thaliana* resistance to *Pst* DC3000/*avrRps4* by increasing its nuclear accumulation and stability

**DOI:** 10.1101/2024.02.29.582862

**Authors:** Baihong Zhang, Shuqin Huang, Shuyu Guo, Yixuan Meng, Yuzhen Tian, Yue Zhou, Hang Chen, Xue Li, Jun Zhou, Wenli Chen

**Author notes:** Corresponding Author: Wenli Chen, e-mail address; Jun Zhou. E-mail address (Baihong Zhang); (Shuqin Huang); (Shuyu Guo); (Yixuan Meng); (Yuzhen Tian); (Yue Zhou); (Hang Chen); (Xue Li).

## Abstract

Autophagy-related gene 6 (ATG6) plays a crucial role in plant immunity. Nonexpressor of pathogenesis-related genes1 (NPR1) acts as a signaling hub of plant immunity. However, the relationship between ATG6 and NPR1 is unclear. Here, we find that ATG6 directly interacts with NPR1. *ATG6* overexpression significantly increased nuclear accumulation of NPR1. Furthermore, we demonstrate that *ATG6* increases NPR1 protein levels and improves its stability. Interestingly, ATG6 promotes the formation of SINCs (SA-induced NPR1 condensates)-like condensates. Additionally, ATG6 and NPR1 synergistically promote the expression of *pathogenesis-related* genes. Further results showed that silencing *ATG6* in *NPR1-GFP* exacerbates *Pst* DC3000/*avrRps4* infection, while double overexpression of *ATG6* and *NPR1* synergistically inhibits *Pst* DC3000/*avrRps4* infection. In summary, our findings unveil an interplay of NPR1 with ATG6 and elucidate important molecular mechanisms for enhancing plant immunity.

**Highlight:** We unveil a novel relationship in which ATG6 positively regulates NPR1 in plant immunity.

## Introduction

Plants are constantly challenged by pathogens in nature. In order to survive and reproduce, plants have evolved complex mechanisms to cope with attack by pathogens (Jones and Dangl, 2006). Nonexpressor of pathogenesis-related genes 1 (NPR1) is a key regulator of plant immunity (Chen *et al*., 2021b). It contains the BTB/POZ (Broad Compex, Tramtrack, and BricaBrac/Pox virus and Zinc finger) domain in the N-terminal region, the ANK (Ankyrin repeats) domain in the middle region, and SA-binding domain (SBD) and the nuclear localization sequence (NLS) in the C-terminal region (Cao *et al*., 1997, Rochon *et al*., 2006, Kumar *et al*., 2022). NPR1 is a receptor of SA (salicylic acid) mainly localized as an oligomer in the cytoplasm and sensitive to the surrounding redox state (Tada *et al*., 2008, Wu *et al*., 2012). SA mediates the dynamic oligomer to dimer response of NPR1 (Tada *et al*., 2008) and promotes translocation of NPR1 into the nucleus, which increases plant resistance to pathogens by activating the expression of immune-related genes (Kinkema *et al*., 2000, Chen *et al*., 2021b).

NPR1 is mainly degraded by the ubiquitin proteasome system (UPS) (Spoel *et al*., 2009, Saleh *et al*., 2015, Skelly *et al*., 2019). An increasing researches have shown that autophagy and the UPS pathway play overlapping roles in regulating intracellular protein homeostasis (Zhou *et al*., 2014, Marshall *et al*., 2015, Kikuchi *et al*., 2020). Our previous study showed that ATGs (autophagy-related genes) are involved in NPR1 turnover (Gong *et al*., 2020). Autophagy negatively regulates *Pst* DC3000/*avrRpm1*-induced programmed cell death (PCD) via the SA receptor NPR1 (Yoshimoto *et al*., 2009). These results imply that ATGs might be involved in plant immunity through the regulation of NPR1 homeostasis. However, the detailed mechanism has not yet been elucidated.

ATG6 is the homologues of yeast Vps30/Atg6 and mammalian BECN1/Beclin1 (Xu *et al*., 2017). It is a common and required subunit of the class III phosphatidylinositol 3-kinase (PtdIns3K) lipid kinase complexes, which regulates autophagosome nucleation in *Arabidopsis thaliana* (*Arabidopsis*) (Qi *et al*., 2017, Wang *et al*., 2020). The homozygous *atg6* mutant is lethal, suggesting that ATG6 is essential for plant growth and development (Fujiki *et al*., 2007, Qin *et al*., 2007, Harrison-Lowe and Olsen, 2008, Patel and Dinesh-Kumar, 2008). In *Arabidopsis*, *N. benthamiana* and wheat, ATG6 or its homologues was reported to act as a positive regulator to enhance plant disease resistance to *P. syringae pv. tomato* (*Pst*) DC3000 and *Pst* DC3000/*avrRpm1* bacteria (Patel and Dinesh-Kumar, 2008), *N. benthamiana* mosaic virus (TMV) (Liu *et al*., 2005), turnip mosaic virus (TuMV) (Li *et al*., 2018), pepper mild mottle virus (PMMoV) (Jiao *et al*., 2020), and *Blumeria graminis f. sp. tritici (Bgt)* fungus (Yue *et al*., 2015). Several research teams have also elucidated that ATG6 interacted with Bax Inhibitor-1 (NbBI-1) (Xu *et al*., 2017) and RNA-dependent RNA polymerase (RdRp) (Li *et al*., 2018) to suppress pathogen infection. However, the mechanism by which ATG6 suppresses pathogen infection by regulating NPR1 has not yet been reported.

Here, we show that ATG6 and NPR1 synergistically enhance *Arabidopsis* resistance to *Pst* DC3000/*avrRps4* infiltration. We discover that ATG6 increases NPR1 protein levels and nuclear accumulation of NPR1. Moreover, ATG6 can stabilize NPR1 and promote the formation of SINCs (SA-induced NPR1 condensates)-like condensates. Our study revealed a unique mechanism in which NPR1 cooperatively increases plant immunity with ATG6.

## Results

### NPR1 physically interacts with ATG6 in vitro and in vivo

To examine the relationship between ATGs and NPRs, we predicted that some ATGs might interact with NPRs. In a yeast two-hybrid (Y2H) screen, we identified that NPR1, NPR3 and NPR4 could interact with ATG6 and several ATG8 isoforms (**Fig. S1** and **Results S1** in the Supplemental Data 1). In this study, we mainly investigated the relationship between ATG6 and NPR1 during the process of plant immune response. Firstly, the NPR1 truncations NPR1-N (1-328AA, containing the BTB/POZ domain, ANK1, ANK2,) and NPR1-C (328-594AA, containing the ANK3, ANK4, SA-binding domain (SBD) and NLS) were used to identify the interaction domains between NPR1 and ATG6. The results showed that NPR1-C interacted with full-length ATG6 in yeast (**Fig. 1a**, line 3). The interaction between NPR1 and SnRK2.8 was used as a positive control (Lee *et al*., 2015). Secondly, pull-down assays were performed *in vitro*. NPR1-His bound GST-ATG6, but not GST (**Fig. 1b**). Furthermore, co-immunoprecipitation (Co-IP) assays were performed in *N. benthamiana*, as shown in **Fig. 1c**, ATG6-mCherry was co-immunoprecipitated with NPR1-GFP. In **Fig. S2**, fluorescence signals of NPR1-GFP and ATG6-mCherry were co-localized in both the nucleus and cytoplasm. The interaction between ATG6 and NPR1 was also verified by a bimolecular fluorescence complementation (BiFC) assay (**Fig. 1d and e**). These results demonstrate that ATG6 interacts with NPR1 both *in vitro* and *in vivo*.

**Figure 1.**
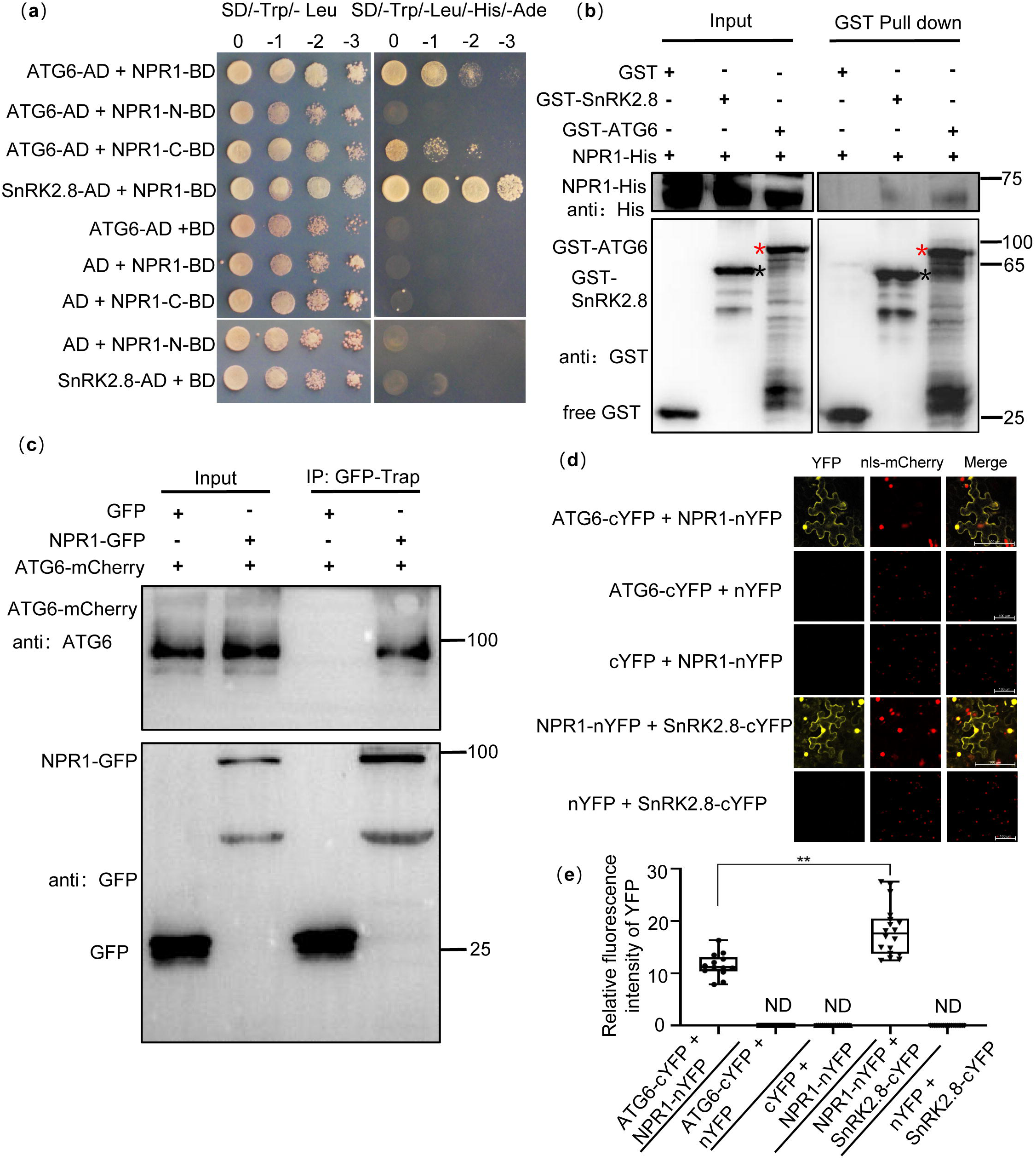
Physical interaction between NPR1 and ATG6. (**a**). Interaction of NPR1 with ATG6 in yeast. The CDS of *ATG6*, *NPR1*, *NPR1-N* (1∼984 bp), *NPR1-C* (984∼1782 bp) and *SnRK2.8* were fused to pGADT7 (AD) and pGBKT7 (BD), respectively. Co-transformation of NPR1-BD + AD, BD + ATG6-AD, BD + SnRK2.8-AD, NPR1-N-BD + AD, NPR1-C-BD + AD were used as negative controls. The interaction of NPR1-BD and SnRK2.8-AD was used as a positive control. Yeast growth on SD/-Trp-Leu-His-Ade media represents interaction. Numbers represent the dilution fold of yeast. 0,-1 (10 fold dilution),-2 (100 fold dilution),-3 (1000 fold dilution). (**b**). In vitro pull-down assays of NPR1-His with GST-ATG6 fusion protein. NPR1-His prokaryotic proteins were incubated with GST-tag Purification Resin conjugated with GST-ATG6, GST and SnRK2.8-GST. Western blotting analysis with anti-GST and anti-His. Black asterisk indicate SnRK2.8-GST bands. Red asterisk indicate GST-ATG6 bands. (**c**). Co-immunoprecipitation of NPR1 with ATG6 in vivo. Total protein was extracted from *N. benthamiana* transiently transformed with ATG6-mCherry + GFP and ATG6-mCherry + NPR1-GFP, followed by IP with GFP-Trap. Western blots analysis with ATG6 and GFP antibodies. (**d**). Bimolecular fluorescence complementation assay of NPR1 with ATG6 in *Nicotiana benthamiana* leaves. Agrobacterium carrying ATG6-nYFP and NPR1-cYFP was co-expressed in leaves of *Nicotiana benthamiana* for 3 days. As a positive control, NPR1-nYFP and SnRK2.8-cYFP were co-expressed. As negative controls, nYFP and ATG6-cYFP, NPR1-nYFP and cYFP, nYFP and SnRK2.8-cYFP were co-expressed. Confocal images were obtained from mCherry, YFP. nls-mCherry as a nuclear localization mark. Scale bar = 100 μm. (**e**). Relative fluorescence intensity of YFP in (d) using image J software, ND means not detected, n = 15 independent images were analyzed to quantify YFP fluorescence. ** indicates that the significant difference compared to the control is at the level of 0.01 (Student t test p value, ** p< 0.01). All experiments were performed with three biological replicates.

### ATG6 co-localized with NPR1 in the nucleus

Remarkably, we found that ATG6 is localized in the cytoplasm and nucleus, and it co-localized with NPR1 in the nucleus (**Fig. S2**). Nuclear localization of ATG6 was also observed in *N. benthamiana* transiently transformed with ATG6-mCherry and ATG6-GFP under normal and SA treatment conditions (**Fig. 2a and b**). ATG6-GFP co-localizes with the nuclear localization marker nls-mCherry (indicated by white arrows) (**Fig. 2b**). Additionally, we observed punctate patterns indicative of canonical autophagy-like localization of ATG6-GFP fluorescence signals (indicated by red circles) (**Fig. 2b**). The nuclear localization signal of ATG6 was also observed in *UBQ10::ATG6-GFP* overexpressing *Arabidopsis* (**Fig. S3a**). To exclude the possibility that the observed localization of ATG6-GFP is due to free GFP. The protein levels of ATG6-GFP and free GFP in *UBQ10::ATG6-GFP Arabidopsis* and *N.benthamiana* were detected before and after SA treatment. Notably, no free GFP was detected and this means that the fluorescence signal observed by confocal microscopy is ATG6-GFP, not free GFP (**Fig. S3d** and **e**). In both plants and animals, proteins are transported to the nucleus via specific nuclear localization signals (NLSs), which are typically characterized by short stretches of basic amino acids (Dingwall and Laskey, 1991, Raikhel, 1992, Nigg, 1997). Furthermore, we analyzed the putative nuclear localization signal (NLS) in the ATG6 protein sequence using NLSExplorer (http://www.csbio.sjtu.edu.cn/bioinf /NLSExplorer). Although we did not identify a classical monopartite NLS, we discovered a bipartite NLS similar to the consensus bipartite sequence (KRX_(10-12)_K(KR)(KR))(Kosugi *et al*., 2009) in the carboxy-terminal region (475-517 aa) of ATG6, with a cut-off score of 2.6 (**Fig. 2c**). Additionally, our comparison of ATG6 C-terminal sequences across several species, including *Microthlaspi erraticum*, *Capsella rubella*, *Brassica carinata*, *Camelina sativa*, *Theobroma cacao*, *Brassica rapa*, *Eutrema salsugineum*, *Raphanus sativus*, *Hirschfeldia incana* and *Brassica napus*, sequence comparison indicates that this bipartite NLS is relatively conserved (**Fig. 2c**).

**Figure 2.**
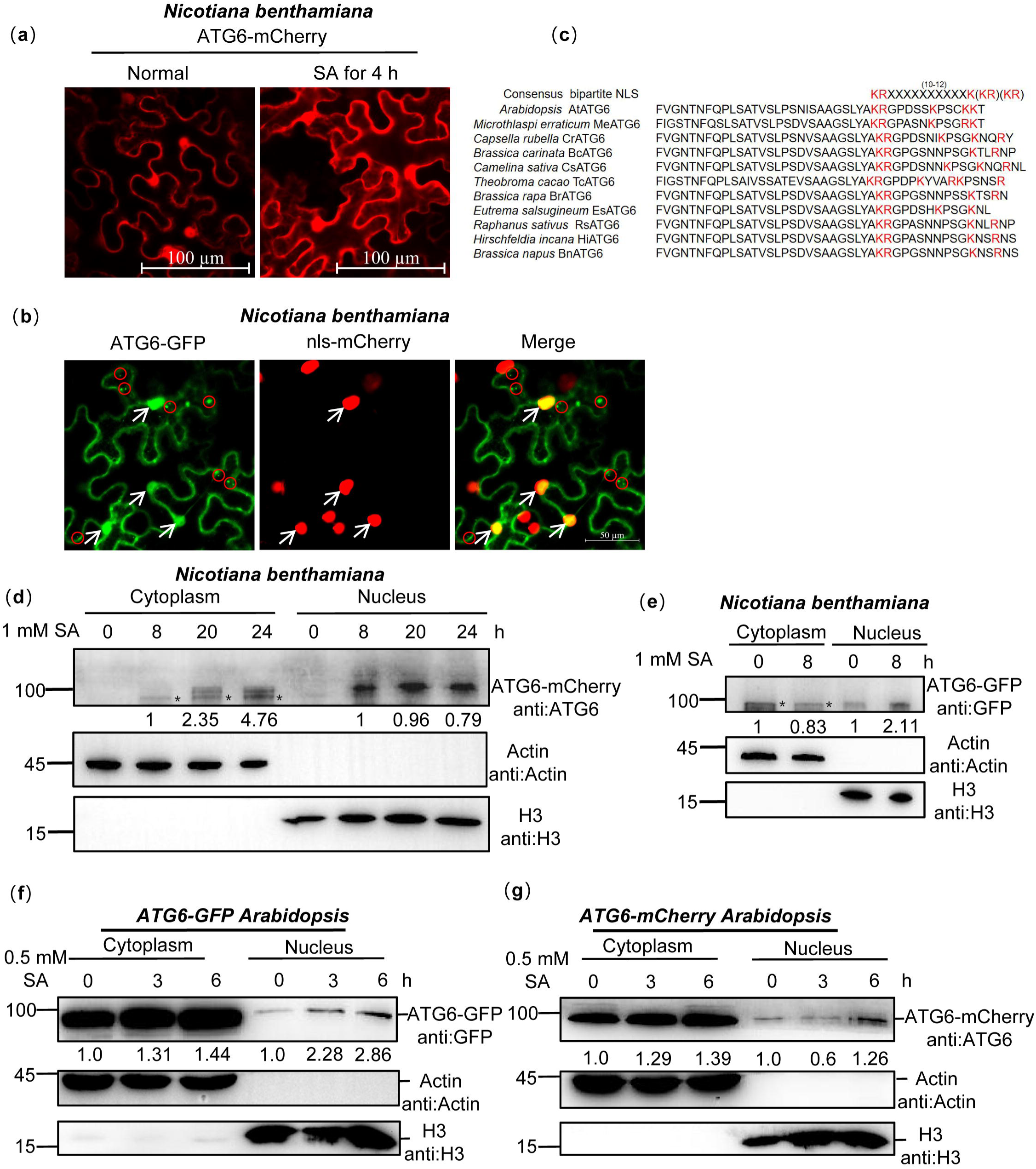
ATG6 is localized in the cytoplasm and nucleus. (**a**). The nuclear localization of ATG6-mCherry in *N. benthamiana*. Scale bar, 100 μm. (**b**). Co-localization of ATG6-GFP and nls-mCherry in *N. benthamiana*. Scale bar, 50 μm. (**c**). ATGs protein nuclear localization sequence analysis using the online NLSExplorer prediction software and sequence comparison of ATG6 C-terminal with other species. (**d**). Subcellular fractionation of ATG6-mCherry in *N. benthamiana* after 1 mM SA treatment. Black asterisk (*) indicate ATG6-mCherry bands. (**e**). Subcellular fractionation of ATG6-GFP in *N. benthamiana* after 1 mM SA treatment. Black asterisk (*) indicate ATG6-GFP bands. (**f**). Subcellular fractionation of ATG6-GFP in *ATG6-GFP Arabidopsis* after 0.5 mM SA treatment. (**g**). Subcellular fractionation of ATG6-mCherry in *ATG6- mCherry Arabidopsis* after 0.5 mM SA treatment. In (**d**-**g**), ATG6-mCherry (**d** and **g**) and ATG6-GFP (**e** and **f**) were detected using ATG6 or GFP antibody. Actin and H3 were used as cytoplasmic and nucleus internal reference, respectively. Numerical values indicate quantitative analysis of ATG6-mCherry and ATG6-GFP using image J. All experiments were performed with three biological replicates.

Moreover, the nuclear and cytoplasmic fractions were separated. Under SA treatment, ATG6-mCherry and ATG6-GFP were detected in the cytoplasmic and nuclear fractions in *N. benthamiana* (**Fig. 2d and e**). However, in *N. benthamiana*, we observed that ATG6-mCherry was not detected in the nuclear fractions under normal conditions, which differents with the results shown in **Fig. 2a**. We suspect that this discrepancy may be due to the fluorescence signal in Fig. 2a primarily arising from free mCherry rather than the ATG6-mCherry fusion. ATG6 was also detected in the nuclear fraction of *UBQ10::ATG6-GFP* and *UBQ10::ATG6-mCherry* overexpressing plants, and SA promoted both cytoplasm and nuclear accumulation of ATG6 (**Fig. 2f and g**). Additionally, we obtained *ATG6* and *NPR1* double overexpression of *Arabidopsis UBQ10::ATG6-mCherry* × *35S::NPR1-GFP* (*ATG6-mCherry* ×*NPR1-GFP*) by crossing and screening (**Fig. S4a**). In *ATG6-mCherry* × *NPR1-GFP*, we observed co-localization of ATG6-mCherry with NPR1-GFP in the nucleus (**Fig. S3b**). These results are consistent with the prediction of the subcellular location of ATG6 in the *Arabidopsis* subcellular database (https://suba.live/) (**Fig. S3c**). Additionally, we have conducted an investigation into the localization of endogenous ATG6 in Col. Our results demonstrate that endogenous ATG6 is present in both the nucleus and cytoplasm, and we have observed that SA treatment promotes the accumulation of ATG6 in the nucleus (**Fig. S5**). Together, these findings suggest that ATG6 is localized to both cytoplasm and nucleus, and co-localized with NPR1 in the nucleus.

### ATG6 overexpression increased nuclear accumulation of NPR1

Previous studies have shown that the nuclear localization of NPR1 is essential for improving plant immunity (Kinkema *et al*., 2000, Chen *et al*., 2021b). We observed that a stronger nuclear localization signal of NPR1-GFP in *ATG6-mCherry* × *NPR1-GFP* leaves than that in *NPR1-GFP* under normal condition and 0.5 mM SA treatment for 3 h (**Fig. 3a-b** and **Fig. S6**). These findings indicate that ATG6 might increase nuclear accumulation of NPR1. To exclude the possibility that the observed localization of NPR1-GFP is due to free GFP, we detected the levels of NPR1-GFP and free GFP in *ATG6-mCherry* x *NPR1-GFP* plants before and after SA treatment. Only ∼ 10 % of free GFP was detected in *ATG6-mCherry* x *NPR1-GFP* plants before and after SA treatment, confirming that the observed localization of NPR1-GFP is not due to free GFP (**Fig. S4b**). Furthermore, the nuclear and cytoplasmic fractions of *ATG6-mCherry* × *NPR1-GFP* and *NPR1-GFP* were separated. Under normal conditions, the nuclear fractions NPR1-GFP in *ATG6-mCherry × NPR1-GFP* and *NPR1-GFP* were relatively weaker (**Fig. 3c**), which differs from the above observation (**Fig. 3a**). We speculate that this phenomenon might be attributed to the rapid turnover of NPR1 in the nucleus (Spoel *et al*., 2009, Saleh *et al*., 2015). Consistent with the fluorescence distribution results, the nuclear fractions of NPR1-GFP in *ATG6-mCherry × NPR1-GFP* were significantly higher than those in *NPR1-GFP* under 0.5 mM SA treatment for 3 and 6 h (**Fig. 3c** and **Fig. S7**). Furthermore, *Agrobacterium* harboring ATG6-mCherry and NPR1-GFP were transiently transformed to *N. benthamiana* leaves. After 1 d, the leaves were treated with 1 mM SA for 8 and 20 h. Subsequently nucleoplasmic separation experiments were performed. Similar to *Arabidopsis*, increased nuclear accumulation of NPR1 was found when *ATG6* was overexpressed (**Fig. 3e** and **Fig. S7**). Notably, we found that the ratio (nucleus NPR1/total NPR1) in *ATG6-mCherry × NPR1-GFP* was not significantly different from that in *NPR1-GFP* after SA treatment, and a similar phenomenon was observed in *N. benthamiana* (**Fig. 3d, 3f** and **Fig. S7**). These results suggested that the increased nuclear accumulation of NPR1 in *ATG6-mCherry* x *NPR1-GFP* plants might attributed to higher levels and more stable NPR1 rather than the enhanced nuclear translocation of NPR1 facilitated by ATG6. Furthermore, we validated the functionality of the ATG6-GFP and ATG6-mCherry fusion proteins utilized in this study by examining the phenotypes of *ATG6-GFP* and *ATG6-mCherry Arabidopsis* plants under carbon starvation conditions (**Fig. S8** and **Results S2** in the Supplemental Data 1).

**Figure 3.**
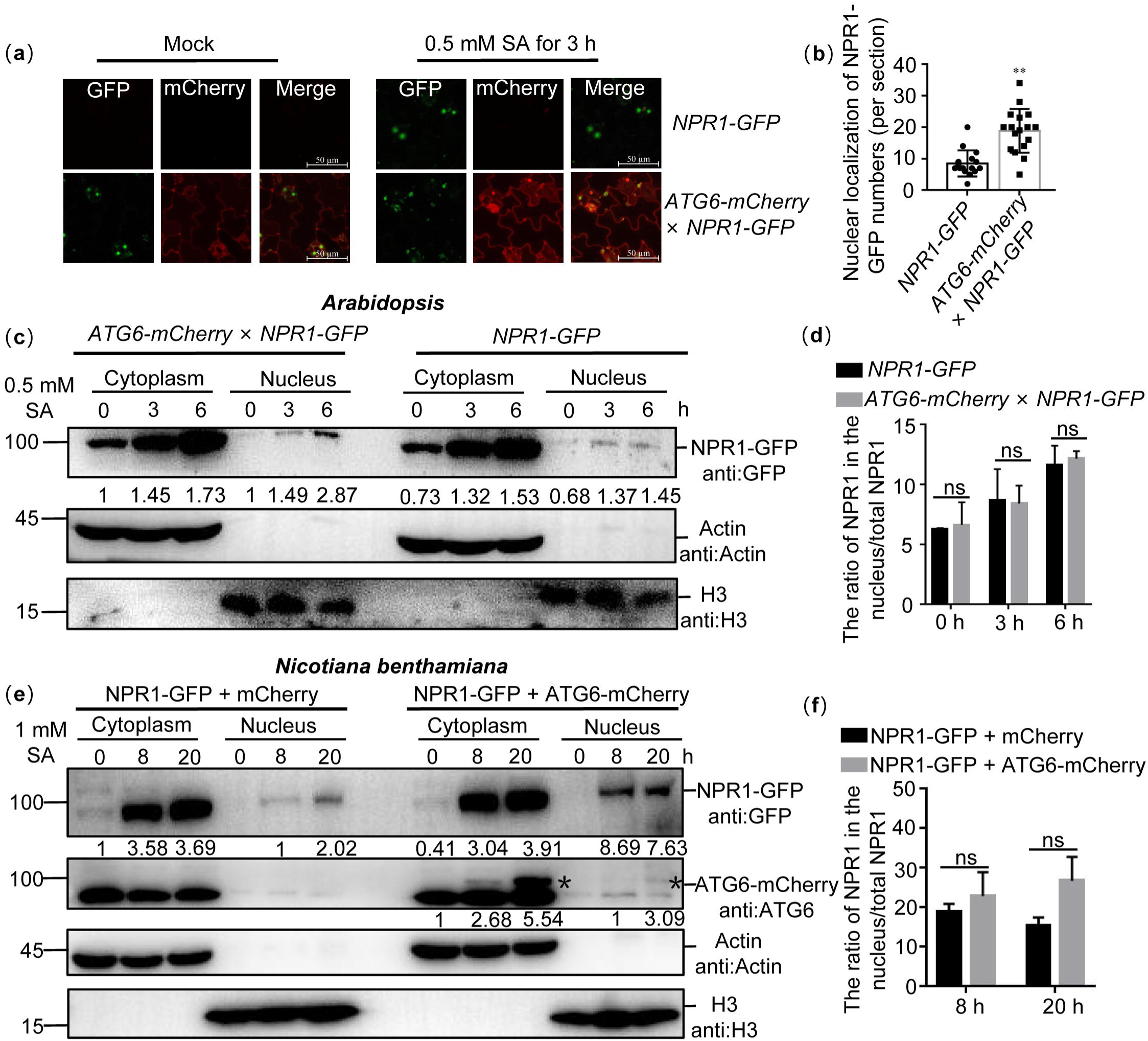
ATG6 increases the nuclear accumulation of NPR1 under SA treatment. (**a**). Confocal images of NPR1-GFP nuclear localization in 7-day-old seedlings of *NPR1-GFP* and *ATG6-mCherry* × *NPR1-GFP* under normal and 0.5 mM SA spray for 3 h. Scale bar, 50 μm. (**b**). The count of nuclear localizations of NPR1-GFP in *ATG6-mCherry × NPR1-GFP* and *NPR1-GFP* Arabidopsis plants following SA treatment. in (**a**), Statistical data were obtained from three independent experiments, each comprising five individual images, resulting in a total of 15 images analyzed for this comparison. ** indicates that the significant difference compared to the control is at the level of 0.01 (Student *t* test *p* value, ** *p*< 0.01). (**c**). Subcellular fractionation of NPR1-GFP in 7-day-old seedlings of *NPR1-GFP* and *ATG6-mCherry* × *NPR1-GFP* after 0.5 mM SA treatment for 0, 3 and 6 h. (**d**). The ration of NPR1 in the nucleus/total NPR1 in (**c**), student’s t-test was conducted to analyze the data. The mean and standard deviation were calculated from three biological replicates, ns indicates no significant difference. (**e**). Subcellular fractionation of NPR1-GFP in *N. benthamiana* after 1 mM SA treatment for 0, 8 and 20 h. (**f**). The ration of NPR1 in the nucleus/total NPR1 in (**e**), student’s t-test was conducted to analyze the data. The mean and standard deviation were calculated from three biological replicates, ns indicates no significant difference. In (**c** and **e**), cytoplasmic and nuclear proteins were extracted from *Arabidopsis* or *N. benthamiana*. NPR1-GFP were detected using GFP antibody. Actin and H3 were used as cytoplasmic and nucleus internal reference, respectively. Numerical values indicate quantitative analysis of NPR1-GFP using image J. All experiments were performed with three biological replicates.

### ATG6 increases endogenous SA levels and promotes the expression of NPR1 downstream target genes

NPR1 localized in the nucleus is essential for activation of immune gene expression (Kinkema *et al*., 2000, Chen *et al*., 2021b). In our study, we observed that *ATG6* overexpression increased nuclear accumulation of NPR1 (**Fig. 3**) and demonstrated an interaction between ATG6 and NPR1 in the nucleus (**Fig. 1d**). Therefore, we speculate that ATG6 might regulate NPR1 transcriptional activity. Notably, the expression level of *ICS1* in *ATG6-mCherry × NPR1-GFP* seedlings was significantly higher than that in *NPR1-GFP* under normal and SA treatment conditions (**Fig. S9**). Free SA levels in *ATG6-mCherry × NPR1-GFP* were also significantly higher compared to *NPR1-GFP* under *Pst* DC3000/*avrRps4* treatment. While there was no significant difference was observed under normal condition (**Fig. 4a**), this may be related to free SA consumption, as it can be converted to bound SA (Ding and Ding, 2020). In addition, the expression of *PR1* (*pathogenesis-related* gene) and *PR5* in *ATG6-mCherry* × *NPR1-GFP* was significantly higher than that of *NPR1-GFP* under normal and SA treatment conditions (**Fig. 4b and c**). The expression of *PR1* and *PR5* in *ATG6-mCherry* was significantly higher than that of Col under *Pst* DC3000/*avrRps4* treatment (**Fig. S10**). These results support the role of ATG6 in facilitating the expression of NPR1 downstream *PR1* and *PR5* genes.

**Figure 4.**
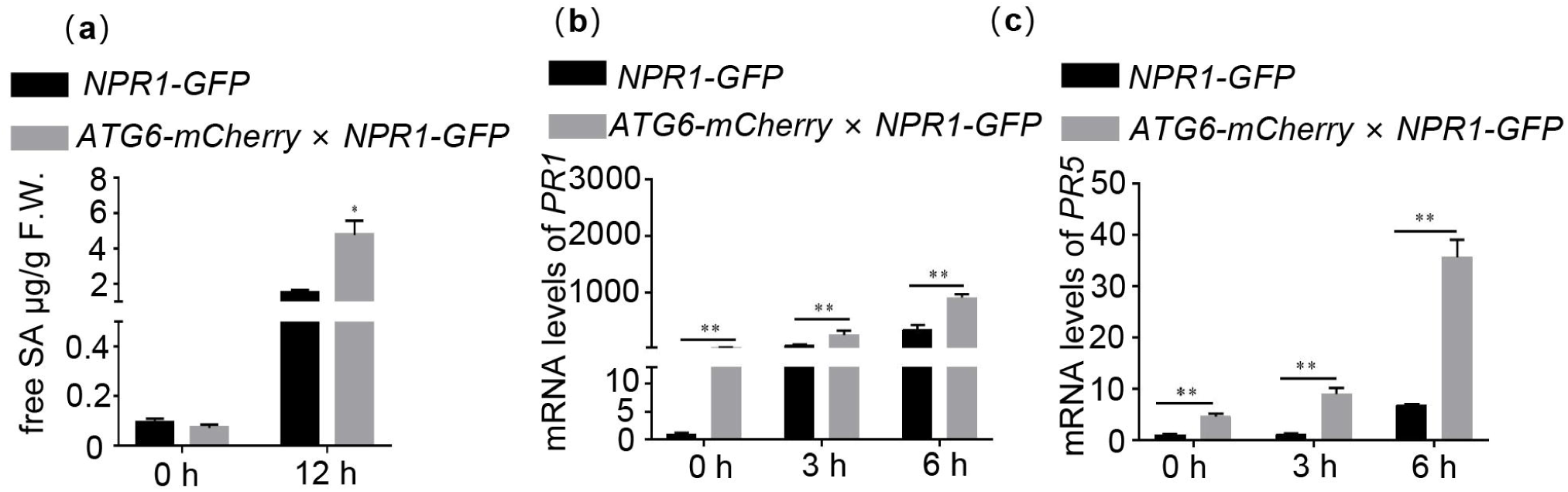
ATG6 increases endogenous SA levels and promotes the expression of NPR1 downstream target genes (a). Level of free SA in 3-week-old *NPR1-GFP* and *ATG6-mCherry* × *NPR1- GFP* after *Pst* DC3000/*avrRps4* for 12 h. (**b-c**). Expression of *PR1* (**b**) and *PR5* (**c**) in 3-week-old *NPR1-GFP* and *ATG6-mCherry* × *NPR1-GFP* under normal and SA treatment conditions, values are means ± SD (n = 3 biological replicates). The *AtActin* gene was used as the internal control. * or ** indicates that the significant difference compared to the control is at the level of 0.05 or 0.01 (Student *t* test *p* value, * *p*< 0.05 or ** *p*< 0.01). All experiments were performed with three biological replicates.

### ATG6 increases NPR1 protein levels and the formation of SINCs-like condensates

Interestingly, similar to previous reports (Zavaliev *et al*., 2020), SA promoted the translocation of NPR1 into the nucleus, but still a significant amount of NPR1 was present in the cytoplasm (**Fig. 3c and e**). Previous studies have shown that SA increased NPR1 protein levels and facilitated the formation of SINCs in the cytoplasm, which are known to promote cell survival (Zavaliev *et al*., 2020). In our experiments, we observed that under SA treatment, the protein levels of NPR1 in *ATG6-mCherry* × *NPR1-GFP* was significantly higher than that in *NPR1-GFP* (**Fig. 5a**). To further support our conclusions, we proceeded to silence *ATG6* in *NPR1-GFP* (*NPR1-GFP*/silencing *ATG6*) and subsequently assessed the protein level of NPR1-GFP before and after SA treatment. Our findings revealed that the protein level of NPR1-GFP in *NPR1-GFP*/silencing *ATG6* under SA treatment was notably lower than that in the *NPR1-GFP*/Negative control (**Fig. S11**). Under SA treatment for 8 h, the protein levels of NPR1-GFP in *N. benthamiana* co-transformed with ATG6- mCherry + NPR1-GFP was also significantly higher than that of mCherry + NPR1-GFP (**Fig. 5b**). While there was a slight increase at 20 h, a minor decrease was observed at 24 h, suggesting that the rise in NPR1 protein levels induced by ATG6 was transient. We also detected the expression of *NPR1* was detected. It is worth noting that NPR1 up-regulation was more obvious in Col after 3 h treatment with *Pst* DC3000/*avrRps4*. After 6 h treatment with *Pst* DC3000/*avrRps4*, there was no significant difference in the expression of NPR1 between Col and *ATG6-mCherry* (**Fig. S12**). These results suggest that ATG6 increases NPR1 protein levels. After SA treatment, more SINCs-like condensates fluorescence were observed in *N. benthamiana* co-transformed with ATG6-mCherry + NPR1-GFP compared to mCherry + NPR1-GFP (**Fig. 5c-d and Supplemental movie 1-2**). Additionally, we observed that SINCs-like condensates signaling partial co-localized with certain ATG6-mCherry autophagosomes fluorescence signals (**Fig. S13**). Taken together, these results suggest that ATG6 increases the protein levels of NPR1 and promotes the formation of SINCs-like condensates, possibly caused by ATG6 increasing SA levels *in vivo*.

**Figure 5.**
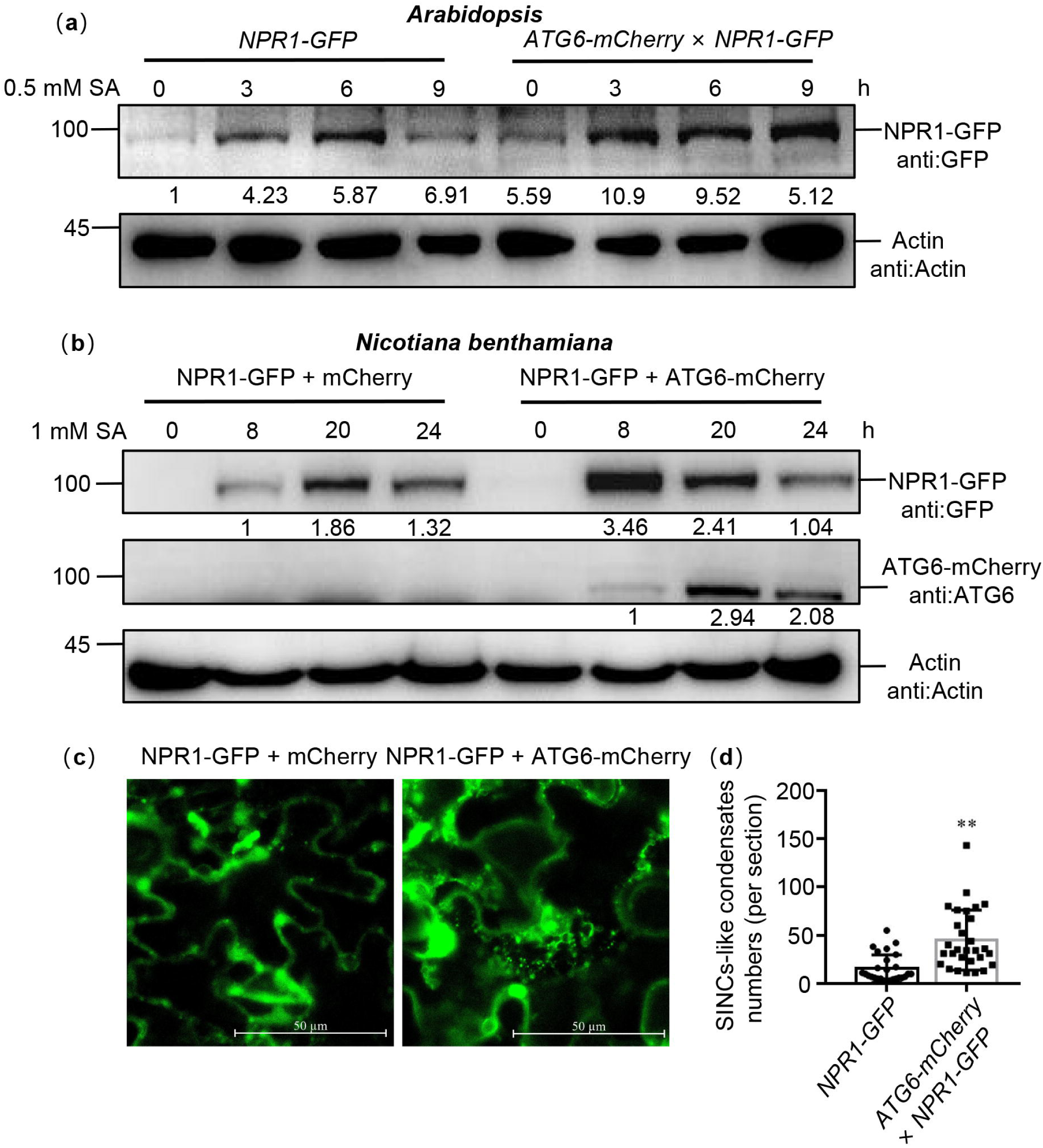
ATG6 increases the NPR1 protein levels and the formation of SINCs-like condensates. (**a**). NPR1-GFP protein levels in 7-day-old seedlings of *NPR1-GFP* and *ATG6-mCherry* × *NPR1-GFP* after 0.5 mM SA treatment for 0, 3, 6 and 9 h. Numerical values indicate quantitative analysis of NPR1-GFP protein using image J. (**b**). NPR1-GFP protein levels in *N. benthamiana*. ATG6-mCherry + NPR1-GFP, NPR1-GFP + mCherry were co-expressed in *N. benthamiana*. After 2 days, leaves were treated with 1 mM SA for 8, 20, 24 h. Total proteins were extracted and analyzed. Numerical values indicate quantitative analysis of NPR1-GFP protein using image J. (**c**). ATG6 promotes the formation of SINCs-like condensates. ATG6-mCherry + NPR1- GFP, NPR1-GFP + mCherry were co-expressed in *N. benthamiana*. After 2 days, leaves were treated with 1 mM SA for 24 h. Confocal images obtained at excitation with wavelengths of 488 nm, scale bar = 50 μm. (**d**). SINCs-like condensates numbers of per section in (**c**), n > 10 sections. ** indicates that the significant difference compared to the control is at the level of 0.01 (Student *t* test *p* value, ** *p*< 0.01). All experiments were performed with three biological replicates.

### ATG6 maintains the protein stability of NPR1

Maintaining the stability of NPR1 is critical for enhancing plant immunity (Skelly *et al*., 2019). To further verify whether ATG6 regulates NPR1 stability, we co-transfected NPR1-GFP with ATG6-mCherry or mCherry in *N. benthamiana* and performed cell-free degradation assays. Our results showed that NPR1-GFP degradation was significantly delayed when *ATG6* was overexpressed (**Fig. S14**). A similar trend was observed in *Arabidopsis*, where the NPR1-GFP protein in *ATG6-mCherry* × *NPR1-GFP* showed a slower degradation rate compared to *NPR1-GFP* during 0∼180 min time period in a cell-free degradation assay (**Fig. 6a and b**). Moreover, when *Arabidopsis* seedlings were treated with cycloheximide (CHX) to block protein synthesis, we found that NPR1-GFP in *NPR1-GFP* was degraded after CHX treatment for 3∼9 h and the half-life of NPR1-GFP is ∼3 h, while the half-life of NPR1- GFP in *ATG6-mCherry* × *NPR1-GFP* is ∼9 h (**Fig. 6c and d**). In addition, we also analyzed the degradation of NPR1-GFP in *NPR1-GFP* and *NPR1- GFP*/*atg5* following 100 μM cycloheximide (CHX) treatment. The results show that the degradation rate of NPR1-GFP in *NPR1-GFP*/*atg5* plants was similarly to that in *NPR1-GFP* plants (**Fig. 6e and f**). These results indicate that ATG6 plays a role in maintaining the stability of NPR1, which may also be related to the fact that ATG6 promotes an increase in free SA *in vivo*, since SA has the function of increasing NPR1 stability (Ding *et al*., 2016, Skelly *et al*., 2019).

**Figure 6.**
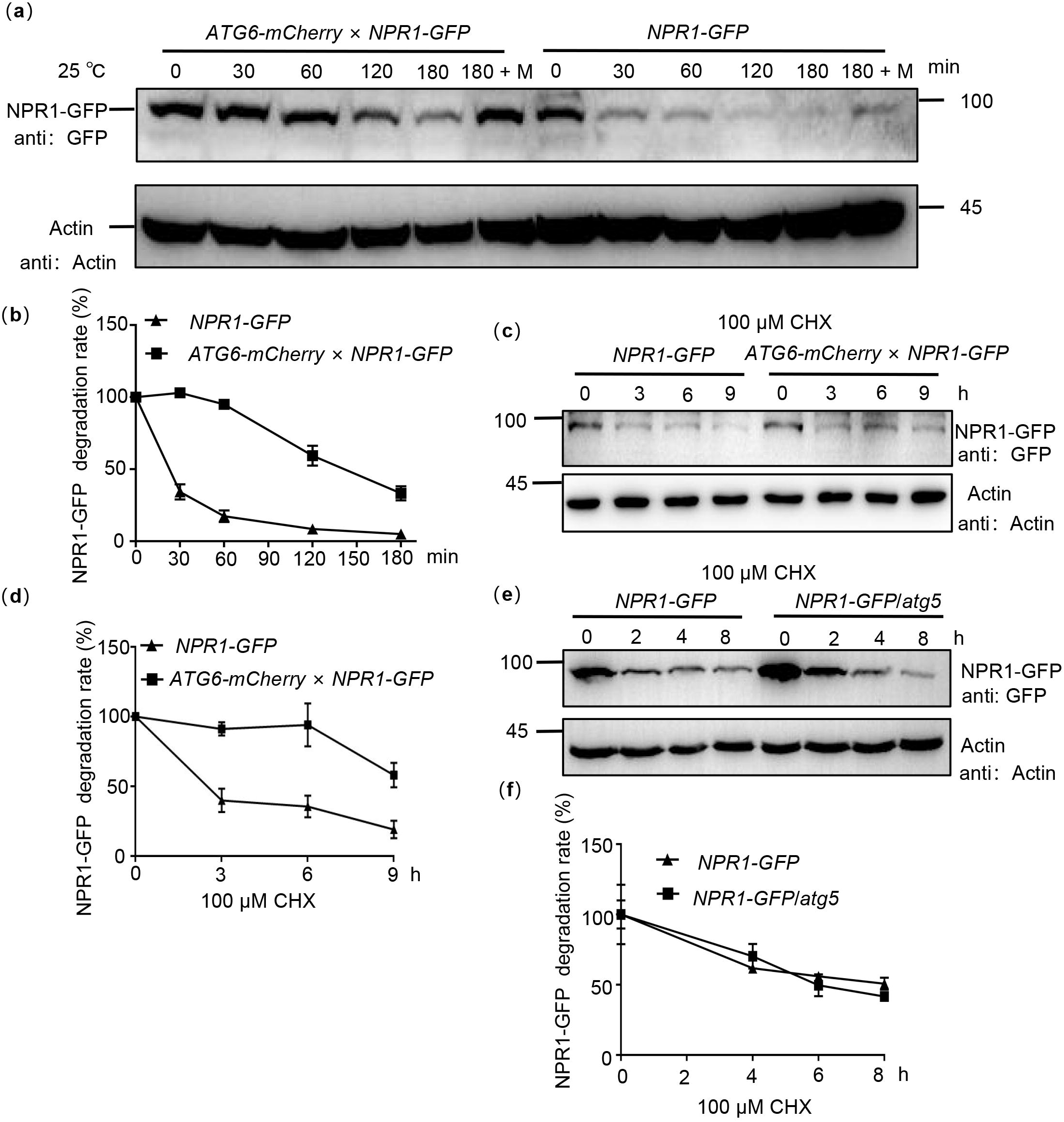
ATG6 improves the protein stability of NPR1. (a). NPR1-GFP degradation assay in *Arabidopsis*. Total proteins from 7-day-old seedlings of *NPR1-GFP* and *ATG6-mCherr*y × *NPR1-GFP* were extracted, using Actin as an internal reference. “M” indicates 100 μM MG115 treatment. (b). Quantification of NPR1-GFP degradation rates in (**a**) using Image J. In (**a** and **b**), the extracts were incubated for 0~180 min at room temperature (25℃), the degradation rate of NPR1-GFP was analyzed. (**c**). NPR1-GFP protein turnover. 7-day-old *NPR1-GFP* and *ATG6-mCherry* × *NPR1-GFP* seedlings were treated with 100 μM cycloheximide (CHX) for different times. Total proteins were analyzed, Actin was used as an internal reference. (**d**). Quantification of NPR1-GFP protein turnover rates in (**c**) using Image J. (**e**). NPR1-GFP protein turnover. 7-day-old *NPR1-GFP* and *NPR1-GFP*/*atg5* seedlings were treated with 100 μM cycloheximide (CHX) for different times. Total proteins were analyzed, Actin was used as an internal reference. (**f**). Quantification of protein levels of NPR1-GFP in (**e**) using Image J. All experiments were performed with three biological replicates.

### ATG6 and NPR1 cooperatively inhibit infection of Pst DC3000/avrRps4

The mRNA expression levels of *ATG6* in Col were significantly increased after 6 h, 12 h and 24 h under *Pst* DC3000/*avrRps4* treatment (**Fig. 7a**). Similarly, both the *ATG6* gene and protein were significantly up-regulated under 0.5 mM SA treatment (**Fig. 7b and c**). These results suggest that the expression of *ATG6* could be induced by *Pst* DC3000/*avrRps4* and 0.5 mM SA treatment.

**Figure 7.**
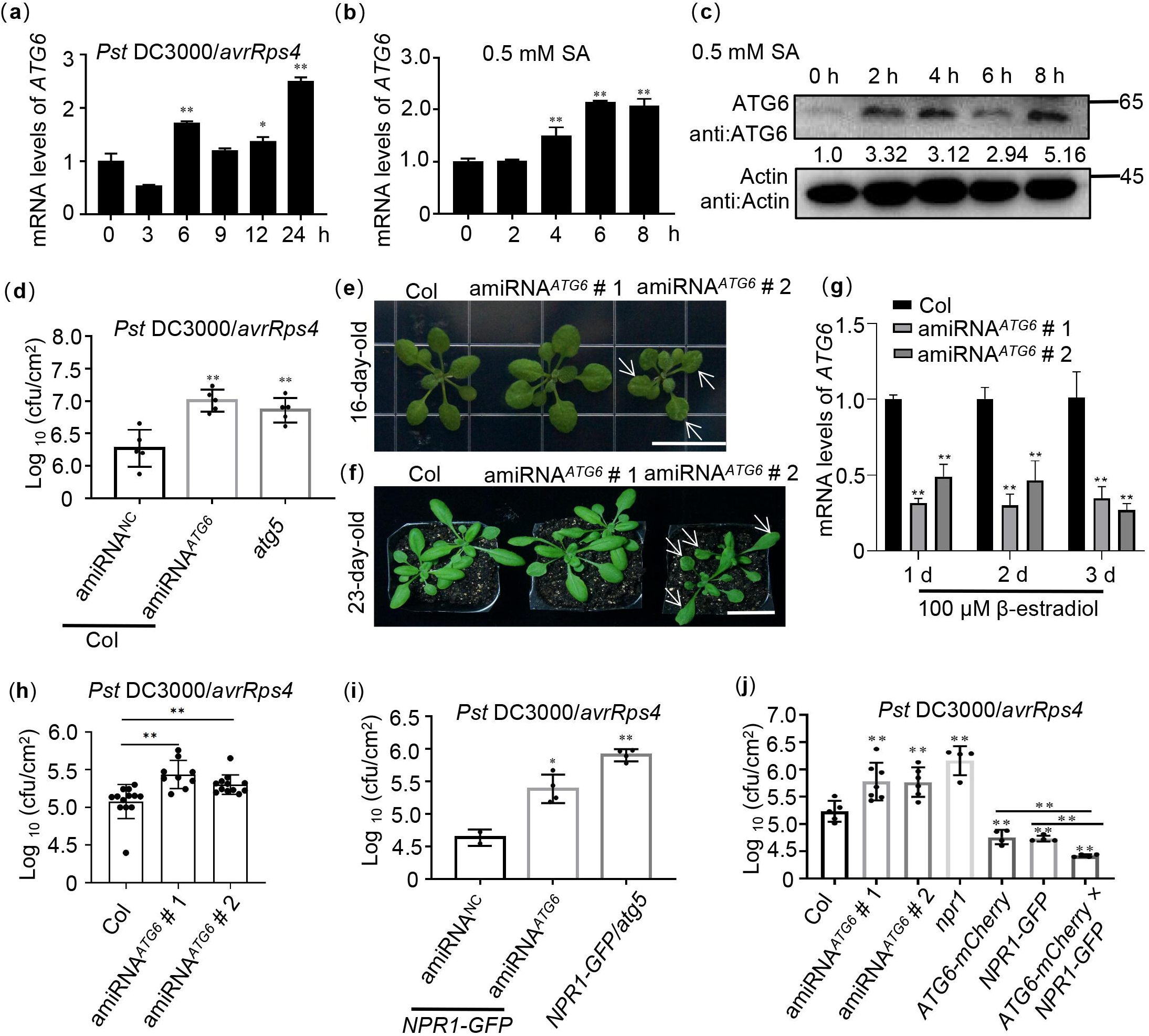
ATG6 and NPR1 cooperatively inhibit the growth of *Pst* DC3000/*avrRps4*. (a). Expression of *ATG6* under *Pst* DC3000/*avrRps4* infiltration in 3-week-old Col leaves, values are means ± SD (n = 3 biological replicates). The *AtActin* gene was used as the internal control. (**b**). Expression of *ATG6* in the presence of 0.5 mM SA in 3-week-old Col leaves, values are means ± SD (n = 3 biological replicates). The *AtActin* gene was used as the internal control. (**c**). The protein levels of ATG6 after 0.5 mM SA in 3-week-old Col leaves. Total leaf proteins from *Arabidopsis* were analyzed, Actin was used as an internal reference. Numerical values indicate quantitative analysis of ATG6 protein using image J. (**d**). Growth of *Pst* DC3000/*avrRps4* in Col/silencing ATG6 and Col/negative control (NC). (**e**). Phenotypes of 16-day-old amiRNA*^ATG6^* # 1 and amiRNA*^ATG6^*# 2. Bar, 1 cm. (**f**). Phenotypes of 23-day-old amiRNA*^ATG6^* # 1 and amiRNA*^ATG6^* # 2. Bar, 3 cm. (**g**). Expression of ATG6 in Col, amiRNA*^ATG6^* # 1 and amiRNA*^ATG6^* # 2 under infiltration treatment of 100 μM β-estradiol, values are means ± SD (n = 3 biological replicates). The *AtActin* gene was used as the internal control. (**h**). Growth of *Pst* DC3000/*avrRps4* in *Arabidopsis* leaves of amiRNA*^ATG6^*# 1, amiRNA*^ATG6^* # 2 and Col. (**i**). Growth of *Pst* DC3000/*avrRps4* in *NPR1-GFP*/silencing *ATG6* and *NPR1-GFP*/NC. (**j**). Growth of *Pst* DC3000/*avrRps4* in *Arabidopsis* leaves of Col, amiRNA*^ATG6^* # 1, amiRNA*^ATG6^* # 2, *npr1*, *NPR1-GFP*, *ATG6-mCherry* and *ATG6-mCherry* × *NPR1-GFP*. In (**d**, **h-j**), a low dose of *Pst* DC3000/*avrRps4* (OD600 = 0.001) was infiltrated. After 3 days, the growth of *Pst* DC3000/*avrRps4* was counted. * or ** indicates that the significant difference compared to the control is at the level of 0.05 or 0.01 (Student *t* test *p* value, * *p*< 0.05 or ** *p*< 0.01). All experiments were performed with three biological replicates.

Considering that ATG6 increases NPR1 protein levels (**Fig. 5a-b**) and promotes its nuclear accumulation (**Fig. 3**), as well as maintains NPR1 stability (**Fig. 6**), then we studied the role of ATG6-NPR1 interactions in plant immunity. However, studying the function of ATG6 is challenging due to the lethality of homozygous *atg6* mutant (Qin *et al*., 2007, Harrison-Lowe and Olsen, 2008, Patel and Dinesh-Kumar, 2008). According to our previous report (Lei *et al*., 2020, Zhang *et al*., 2023), *ATG6* was silenced using artificial miRNA*^ATG6^* (amiRNA*^ATG6^*) delivered by the gold nanoparticles (AuNPs). First, the effect of *ATG6* silencing in Col on the plant immune response was investigated. Similar to *atg5*, Col/silencing *ATG6* exhibited more active growth of *Pst* DC3000/*avrRps4* than Col/negative control (NC) after *Pst* DC3000/*avrRps4* infiltration for 3 days (**Fig. 7d**). Furthermore, according to the previously reported methods (Ohira *et al*., 2017, Gomez *et al*., 2022), we generated two amiRNA*^ATG6^*lines (amiRNA*^ATG6^ #* 1 and amiRNA*^ATG6^ #* 2) designed against *ATG6* and placed under the control of a β-estradiol inducible promoter. There were no significant phenotypic differences in amiRNA*^ATG6^ #* 1 compared to the Col, while amiRNA*^ATG6^ #* 2 exhibited a slight leaf developmental defect (**Fig. 7e and f**). Subsequently, we investigated the expression of *ATG6* following treatment with 100 μM β-estradiol. Our results showed that, after 100 μM β-estradiol treatment for 1∼3 d, the expression of *ATG6* in both amiRNA*^ATG6^*# 1 and amiRNA*^ATG6^* # 2 lines was significantly lower than that in Col. Specifically, the expression of *ATG6* in the amiRNA*^ATG6^ #1* and amiRNA*^ATG6^ #2* lines decreased by 50∼70% compared with Col (**Fig. 7g**). Furthermore, to assess the function of ATG6 in plant immune, we performed infiltrations of *Pst* DC3000/*avrRps4* after 100 uM β-estradiol treatment for 24 h. We compared the growth of *Pst* DC3000/*avrRps4* in the amiRNA*^ATG6^* lines and Col. The results clearly demonstrate that the growth of *Pst* DC3000/*avrRps4* in amiRNA*^ATG6^ #* 1 and amiRNA*^ATG6^ #* 2 was significantly more compared to Col (**Fig. 7h**). Moreover, we silenced *ATG6* in *NPR1-GFP* (*NPR1-GFP*/silencing *ATG6*), and *NPR1-GFP*/*atg5* (crossed *NPR1-GFP* with *atg5* to obtain *NPR1-GFP*/*atg5*) was used as an autophagy-deficient control. There was more *Pst* DC3000/*avrRps4* growth in *NPR1-GFP*/silencing *ATG6* and *NPR1-GFP*/*atg5* compared to *NPR1-GFP*/NC after *Pst* DC3000/*avrRps4* infiltration (**Fig. 7i**). In contrast, the growth of *Pst* DC3000/*avrRps4* in *NPR1- GFP*, *ATG6-mCherry*, *ATG6-mCherry × NPR1-GFP* was significantly lower than that in Col and *npr1* (**Fig. 7j**) and was the lowest in *ATG6-mCherry* × *NPR1-GFP* (**Fig. 7j**).

These results confirm that ATG6 and NPR1 cooperatively enhance *Arabidopsis* resistance to inhibit *Pst* DC3000/*avrRps4* infection. Together, these results suggest that ATG6 improves plant resistance to pathogens by regulating NPR1.

## Discussion

Although SA signaling and autophagy are related to the plant immune system (Yoshimoto *et al*., 2009, Munch *et al*., 2014, Wang *et al*., 2016), the connection of these two processes in plant immune processes and their interaction is rarely reported. Previous studies have shown that unrestricted pathogen-induced PCD requires SA signalling in autophagy-deficient mutants. SA and its analogue benzo (1,2,3) thiadiazole-7-carbothioic acid (BTH) induce autophagosome production (Yoshimoto *et al*., 2009). Moreover, autophagy has been shown to negatively regulates *Pst* DC3000/*avrRpm1*-induced PCD via the SA receptor NPR1 (Yoshimoto *et al*., 2009), implying that autophagy regulates SA signaling through a negative feedback loop to limit immune-related PCD. Here, we demonstrated that ATG6 increases NPR1 protein levels and nuclear accumulation (**Fig. 3** and **5**). Additionally, ATG6 also maintains the stability of NPR1 and promotes the formation of SINCs-like condensates (**Fig. 5** and **6**). These findings introduce a novel perspective on the positive regulation of NPR1 by ATG6, highlighting their synergistic role in enhancing plant resistance.

Our results confirmed that *ATG6* overexpression significantly increased nuclear accumulation of NPR1 (**Fig. 3**). ATG6 also increases NPR1 protein levels and improves NPR1 stability (**Fig. 5 and 6**). Therefore, we consider that the increased nuclear accumulation of NPR1 in *ATG6-mCherry* x *NPR1-GFP* plants might result from higher levels and more stable NPR1 rather than the enhanced nuclear translocation of NPR1 facilitated by ATG6. To verify this possibility, we determined the ratio of NPR1-GFP in the nuclear localization versus total NPR1-GFP. Notably, the ratio (nucleus NPR1/total NPR1) in *ATG6-mCherry* × *NPR1-GFP* was not significantly different from that in *NPR1-GFP*, and there is a similar phenomenon in *N. benthamiana* (**Fig. 3c-f**). Further we analyzed whether ATG6 affects NPR1 protein levels and protein stability. Our results show that ATG6 increases NPR1 protein levels under SA treatment and ATG6 maintains the protein stability of NPR1 (**Fig. 5 and 6**). These results suggested that the increased nuclear accumulation of NPR1 by ATG6 result from higher levels and more stable NPR1.

NPR1 is an important signaling hub of the plant immune response. Nuclear localization of NPR1 is essential to enhance plant resistance (Kinkema *et al*., 2000, Chen *et al*., 2021b), it interacts with transcription factors such as TGAs in the nucleus to activate expression of downstream target genes (Chen *et al*., 2019, Chen *et al*., 2021a). A recent study showed that nuclear-located ATG8h recognizes C1, a geminivirus nuclear protein, and promotes C1 degradation through autophagy to limit viral infiltration in solanaceous plants (Li *et al*., 2020). Here, we confirmed that ATG6 is also distributed in the nucleus and ATG6 is co-localized with NPR1 (**Fig. 1d** and **2**), suggesting that ATG6 interact with NPR1 in the nucleus. ATG6 synergistically inhibits the infection of *Pst* DC3000/*avrRps4* with NPR1. Chen et al. found that in the nucleus, NPR1 can recruit enhanced disease susceptibility 1 (EDS1), a transcriptional coactivator, to synergistically activate expression of downstream target genes (Chen *et al*., 2021a). Previous studies have shown that acidic activation domains (AADs) in transcriptional activators (such as Gal4, Gcn4 and VP16) play important roles in activating downstream target genes. Acidic amino acids and hydrophobic residues are the key structural elements of AAD (Pennica *et al*., 1984, Cress and Triezenberg, 1991, Van Hoy *et al*., 1993). Chen et al. found that EDS1 contains two ADD domains and confirmed that EDS1 is a transcriptional activator with AAD (Chen *et al*., 2021a). Here, we also have similar results that *ATG6* overexpression significantly enhanced the expression of *PR1* and *PR5* (**Fig. 4b-c and Fig.S10**), and that the ADD domain containing acidic and hydrophobic amino acids is also found in ATG6 (148-295 AA) (**Fig. S15**). We speculate that ATG6 might act as a transcriptional coactivator to activate *PRs* expression synergistically with NPR1.

A recent study showed that SA not only enhances plant resistance by increasing NPR1 nuclear import and transcriptional activity, but also promotes cell survival by coordinating the distribution of NPR1 in the nucleus and cytoplasm (Zavaliev *et al*., 2020). Notably, NPR1 accumulated in the cytoplasm recruits other immunomodulators (such as EDS1, PAD4 etc) to form SINCs to promote cell survival (Zavaliev *et al*., 2020). Similarly, we also found that NPR1 accumulated abundantly in the cytoplasm after SA treatment and that ATG6 significantly increased NPR1 protein levels (**Fig. 3c, 3e and 5a-b**). Obviously, the accumulation of NPR1 in the cytoplasm may be related to ATG6 synergizing with NPR1 to enhance plant resistance. Interestingly, *ATG6* overexpression significantly increased the formation of SINCs-like condensates (**Fig. 5c-d** and **Supplemental movie 1-2**), which should also be a way for ATG6 and NPR1 to synergistically resist infection of pathogens. We consider that ATG6 promotes the formation of SINCs-like condensates through the dual action of endogenous and exogenous SA. Considering that ATG6 promotes SINCs-like condensates formation, we further examined changes in cell death in Col, amiRNA*^ATG6^*# 1, amiRNA*^ATG6^* # 2, *npr1*, *NPR1- GFP*, *ATG6-mCherry* and *ATG6-mCherry* × *NPR1-GFP* plants. The results of Taipan blue staining showed that *Pst* DC3000/*avrRps4-*induced cell death in *npr1*, amiRNA*^ATG6^ #* 1 and amiRNA*^ATG6^ #* 2 was significantly higher compared to Col (**Fig. S16**). Conversely, *Pst* DC3000/*avrRps4-*induced cell death in *ATG6-mCherry*, *NPR1-GFP* and *ATG6-mCherry × NPR1-GFP* was significantly lower compared to Col. Notably, *Pst* DC3000/*avrRps4-*induced cell death in *ATG6-mCherry × NPR1-GFP* was significantly lower compared *ATG6-mCherry* and *NPR1-GFP* (**Fig. S16**). These results suggest that ATG6 and NPR1 cooperatively inhibit *Pst* DC3000/*avrRps4*-induced cell dead.

ATG6 is a common and required subunit of PtdIns3K lipid kinase complexes, which regulates autophagosome nucleation in *Arabidopsis* (Qi *et al*., 2017, Bozhkov, 2018). In this study, we also found that ATG6 can maintain the stability of NPR1. Thus, to confirm whether the regulation of NPR1 protein stability by ATG6 is autophagy-dependent, we used autophagy inhibitors (Concanamycin A, ConA and Wortmannin, WM) to detect the degradation of NPR1-GFP. Cell-free degradation assays showed that 100 μM MG115 treatment significantly inhibited the degradation of NPR1-GFP. However, 5 μM concanamycin A treatment did not significantly delay NPR1 degradation (**Fig. S17**). Remarkably, treatment with 30 μM Wortmannin resulted in a slight acceleration of NPR1 degradation, while the combined treatment of ConA and WM significantly expedited the degradation of NPR1 (**Fig. S17**). This may be related to crosstalk between autophagy and 26S Proteasome. It has been demonstrated that autophagy directly regulates the activity of the 26S proteasome under normal conditions or treatment with *Pst* DC3000 (Marshall *et al*., 2015, Ustun *et al*., 2018). Marshall et al. found that the 26S proteasome subunits (RPN1, RPN3, RPN5, RPN10, PAG1, PBF1) are significantly enriched in autophagy-deficient mutantsunder normal growth conditions (Marshall *et al*., 2015). Treatment with concanamycin A (ConA), an inhibitor of vacuolar-type ATPase, increased the level of the 20S proteasome subunit PBA1 under treatment with *Pst* DC3000 (Ustun *et al*., 2018). In addition, we also analyzed the degradation of NPR1-GFP in *NPR1-GFP* and *NPR1-GFP*/*atg5* following 100 μM cycloheximide (CHX) treatment. The results show that the degradation rate of NPR1-GFP in *NPR1-GFP*/*atg5* plants was similarly to that in *NPR1-GFP* plants (**Fig. 6e and f**). These results suggest that deletion of ATG5 do not affect the protein stability of NPR1.

An increasing number of studies have shown that ATGs differentially affect plant immunity. Deletion of ATGs (ATG5, ATG7, ATG10 etc.) leads to reduced resistance of plants to necrotrophic pathogens (Lai *et al*., 2011, Lenz *et al*., 2011, Minina *et al*., 2018). ATGs can directly interact with other proteins to positively regulate plant immunity. In *N. benthamiana*, ATG8f interacts the effector protein βC1 of the cotton *leaf curl multan virus* and promotes its degradation to limit pathogen infection (Haxim *et al*., 2017). Notably, ATG18a can interact with WRKY33 transcription factor to synergistically against *Botrytis* infection (Lai *et al*., 2011). Our evidence shows that ATG6 interacts with NPR1 and works together to counteract pathogen infection by positively regulating NPR1 and SA levels *in vivo*. In conclusion, we unveil a novel relationship in which ATG6 positively regulates NPR1 in plant immunity (**Fig. 8**). ATG6 interacts with NPR1 to synergistically enhance plant resistance by regulating NPR1 protein levels, stability, nuclear accumulation, and formation of SINCs-like condensates.

**Figure 8.**
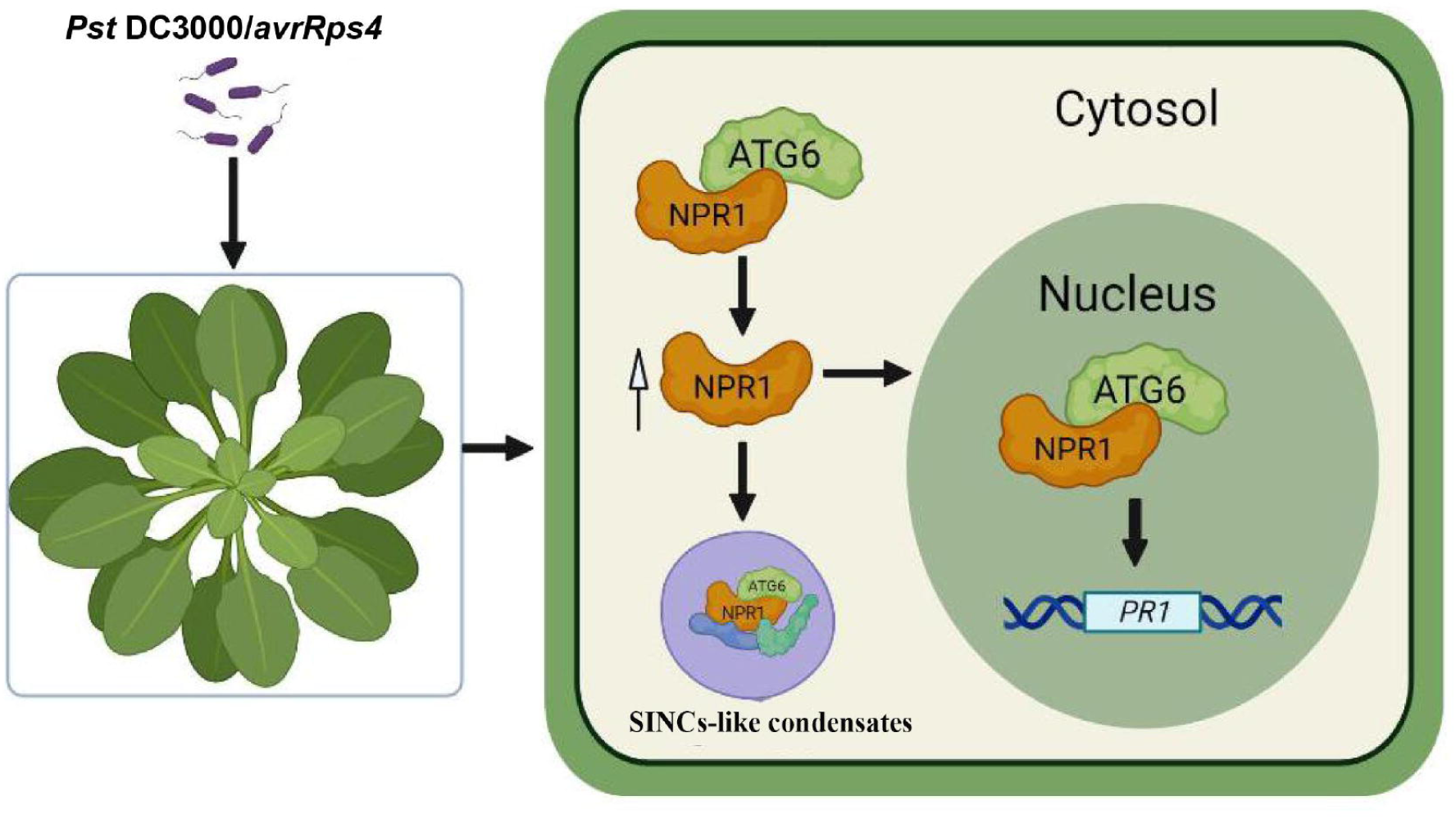
Working model for NPR1 regulation by ATG6. ATG6 interacts directly with NPR1 to increase NPR1 protein level and stability, thereby promoting the formation of SINCs-like condensates and increasing the nuclear accumulation of NPR1. ATG6 synergistically activates *PRs* expression with NPR1 to cooperatively enhance resistance to inhibit *Pst* DC3000/*avrRps4* infection in *Arabidopsis*.

## Material and Methods

### Plasmid construction

Details of plasmid construction methods are listed in Methods S1 of Supplemental Data 1, primer used are listed in Table S1 and S2 of Supplemental Data 1. The mapping of vectors is listed in Supplemental Data 2.

### Plant material Arabidopsis

*35S::NPR1-GFP* (in *npr1-2* background) and *npr1-1* were kindly provided by Dr. Xinnian Dong of Duke University; *atg5-1* (SALK_020601).

*UBQ10::ATG6-mCherry*, *UBQ10::ATG6-GFP and* amiRNA*^ATG6^* lines were obtained by *Agrobacterium* transformation (Clough and Bent, 1998). *ATG6*, *NPR1* double overexpression of *Arabidopsis* (*ATG6-mCherry* × *NPR1-GFP*) and *NPR1-GFP*/*atg5* were obtained by crossing respectively.

Full description of the *Arabidopsis* screening is included Methods S2 of Supplemental Data 1. Details of plant material are listed in Table S3 of Supplemental Data 1.

### Growth conditions Arabidopsis thaliana

All *Arabidopsis thaliana* (*Arabidopsis*) seeds were treated in 10% sodium hypochlorite for 7 min, washed with ddH_2_O, and treated in 75% ethanol for 30 s, finally washed three times with ddH_2_O. Seeds were sown in 1/2 MS with 2% sucrose solid medium, vernalized at 4℃ for 2 days.

For 7-day-old *Arabidopsis* seedling cultures, the plates were placed under the following conditions: daily cycle of 16 h light (∼80 µmolm^−2^. s^−1^) and 8 h dark at 23 ± 2℃.

For 3-week-old *Arabidopsis* cultures, after 7 days of growth on the plates, the seedlings were transferred to soil for further growth for 2 weeks under the same conditions (Zhang *et al*., 2018).

### N. benthamiana

For 3-week-old *N. benthamiana* cultures, seeds were sown in the soil and vernalized at 4℃ for 2 days. After 10 days of growth on soil, the seedlings were transferred to soil for further growth for 2 weeks under the same conditions (Jiao *et al*., 2019).

### Treatment conditions

#### Treatment of 7-day-old seedlings

##### For SA treatment

7-day-old *Arabidopsis* seedlings were transferred to 1/2 MS liquid medium containing 0.5 mM SA for 0, 3 and 6 h, respectively. The corresponding results are shown in Fig. 2f, g, 3c, d and 5a.

##### For cycloheximide (CHX) treatment

Seedlings of *Arabidopsis* (7 days) were transferred to 1/2 MS liquid medium containing 100 μM CHX for 0, 3, 6 and 9 h, respectively. The corresponding results are shown in Fig. 6c and e.

#### Treatment of 3-week-old Arabidopsis

##### For silencing ATG6 in Col and NPR1-GFP

As previously described (Lei *et al*., 2020, Zhang *et al*., 2022, Zhang *et al*., 2023), 1 mM gold nanoparticles (AuNPs) were synthesized. The artificial microRNA (amiRNA)*^ATG6^* (UCAAUUCUAGGAUAACUGCCC) was designed based on the Web MicroRNA Designer (http://wmd3.weigelworld.org/) platform. The complementary sequence of amiRNA*^ATG6^* is located on the eighth exon of the *ATG6* gene. The sequence of “UUCUCCGAACGUGUCACGUTT” was used as a negative control (NC). NC is a universal negative control without species specificity (Gao *et al*., 2018, Lei *et al*., 2020). amiRNA*^ATG6^* and amiRNA^NC^ synthesized by Suzhou GenePharma. AuNPs (1 mM) and amiRNA*^ATG6^* (20 µM) were incubated at a 9:1 ratio for 30 min at 25℃, 50 rpm. After incubation, a mixture of AuNPs and amiRNA*^ATG6^* was diluted 15-fold with the infiltration buffer (pH 5.7, 10 mM MES, 10 mM MgCl_2_) and infiltrated through the abaxial leaf surface into 3- week-old Col or *NPR1-GFP* for 1-3 days. The third day was chosen as material for *ATG6* silencing. After the third day of AuNPs-amiRNA*^ATG6^*and AuNPs-amiRNA^NC^ infiltration, *Pst* DC3000/*avrRps4* was infiltrated, and then growth of *Pst* DC3000/*avrRps4* was detected.

##### For β-estradiol treatment

100 μM β-estradiol was infiltrated to treat 3-week-old *Arabidopsis* leaves. After 24 h of treatment with β-estradiol, *Pst* DC3000/*avrRps4* was infiltrated and then growth of *Pst* DC3000/*avrRps4* was detected after 3 d.

##### For Pst DC3000/avrRps4 infiltration

Infiltration with *Pst* DC3000/*avrRps4* was performed as previously described (Wang *et al*., 2016, Skelly *et al*., 2019). Full description of the *Pst* DC3000/*avrRps4* culture is included in Methods S3 of Supplemental Data 1.

##### For SA treatment

For 3-week-old Col, 0.5 mM SA was infiltrated into the leaves for 0, 2, 4, 6 and 8 h. The corresponding results are shown in Fig. 7b and c.

#### Yeast two-hybrid assay

Yeast two-hybrid experiments were performed according to the previously described protocol (Fu *et al*., 2012). Full description of Yeast two-hybrid is included in Methods S4 of Supplemental Data 1.

#### Pull down assays in vitro

500 μL of GST, GST-ATG6, SnRK2.8-GST were incubated with GST-tag Purification Resin (Beyotime, P2250) for 2 h at 4°C. The mixture was then centrifuged at 1500 g for 1 min at 4°C, and the resin was washed three times with PBS buffer. Next, the GST-tag purification resin was incubated with the NPR1-His for 2 h at 4°C. After washing three times with PBS buffer, 2 × sample buffers were added to the resin and denatured at 100 °C for 10 mins. The resulting samples were then used for western blotting analysis. Full description of prokaryotic proteins expression is included in Methods S5 of Supplemental Data 1.

#### Co-immunoprecipitation

0.5 g leaves of *N. benthamiana* transiently transformed with ATG6-mCherry + GFP and ATG6-mCherry + NPR1-GFP were fully ground in liquid nitrogen and homogenized in 500 μL of lysis buffer (50 mM Tris-HCl pH 7.5, 150 mM NaCl, 0.5 mM EDTA, 5% Glycerol, 0.2% NP40, 1 mM PMSF, 40 μM MG115, protease inhibitor cocktail 500× and phosphatase inhibitor cocktail 5000×). The samples were then incubated on ice for 30 mins, and centrifuged at 10142 g (TGL16, cence, hunan, China) for 15 mins at 4°C. The supernatant (500 μL) was incubated with 20 μL of GFP-Trap Magnetic Agarose beads (ChromoTek, gtma-20) in a 1.5 mL Eppendorf tube for 2 h by rotating at 4°C. After incubation, the GFP-Trap magnetic Agarose beads were washed three times with cold wash buffer (50 mM Tris-HCl pH 7.5, 150 mM NaCl, 0.5 mM EDTA) and denatured at 75°C for 10 minutes after adding 2 × sample buffer. Western blotting was performed with antibodies to ATG6 and GFP.

#### Nuclear and cytoplasmic separation

Nuclear and cytoplasmic separation were performed according to the previously described method (Kinkema *et al*., 2000). Full description of nuclear and cytoplasmic separation is given in Methods S6 of Supplemental Data 1.

#### Protein degradation in vitro

Protein degradation assays were performed according to a previously described method (Spoel *et al*., 2009, Saleh *et al*., 2015). Full description of protein degradation is included in Methods S7 of Supplemental Data 1.

#### Protein Extraction and Western Blotting Analysis

Protein extraction and western blotting were performed as previously described (Lei *et al*., 2020, Zhang *et al*., 2022). Protein was denatured at 100°C for 10 mins. NPR1 protein was denatured at 75°C for 10 mins (Lei *et al*., 2020). Full description is included in Methods S8 of Supplemental Data 1. Antibody information is presented in Table S4 of Supplemental Data 1.

### Confocal microscope observation

#### For nuclear localization of NPR1-GFP observation

7-day-old seedlings of *NPR1-GFP* and *ATG6-mCherry × NPR1-GFP* were sprayed with 0.5 mM SA for 0 and 3 h. GFP and mCherry fluorescence signals in leaves were observed under the confocal microscope (Zeiss LSM880). Statistical data were obtained from three independent experiments, each comprising five individual images, resulting in a total of 15 images analyzed for this comparison.

#### For the Bimolecular Bluorescence Complementation assay (BiFC)

*Agrobacterium* was infiltrated into *N. benthamiana* as previously described (Jiao *et al*., 2019). Fluorescence signals were observed after 3 days. The full description of BiFC is contained in Methods S9 and S10 of Supplemental Data 1.

#### For the observation of SINCs-like condensates

*Agrobacterium* was infiltrated into *N. benthamiana*. After 2 days, the leaves were treated in 1 mM SA solution for 24 h, and then fluorescence signals were observed. At least 20 image sets were obtained and analyzed. A full description of SINCs-like condensates observation is included in Methods S11 of Supplemental Data 1.

#### For growth of Pst DC3000/avrRps4

A low dose (OD_600_ = 0.001) of *Pst* DC3000/*avrRps4* was used for the infiltration experiments. After 3 days, the colony count was counted according to a previous description (Wang *et al*., 2016, Lei *et al*., 2020). Full description of the growth of *Pst* DC3000/*avrRps4* is given in Methods S12 of Supplemental Data 1.

#### Free SA measurement

Free SA was extracted from 3-week-old *Arabidopsis* using a previously described method (Wang *et al*., 2016, Gong *et al*., 2020). Free SA was measured by High-performance liquid chromatography (Shimadzu LC-6A, Japan). Detection conditions: 294 nm excitation wavelength, 426 nm emission wavelength.

#### Real-Time Quantitative PCR (RT-qPCR)

Total RNA was extracted from *Arabidopsis* (100 mg) using Trizol RNA reagent (Invitrogen, 10296-028, Waltham, MA, USA). RT-qPCR assays were performed as previously described (Zhang *et al*., 2018, Zhang *et al*., 2022). All primers for RT-qPCR are listed individually in Table S5 of Supplemental Data 1. Full description of RT-qPCR is included in Methods S13 of Supplemental Data 1.

#### Trypan Blue Staining

The leaves of 3-week-old Col, amiRNA*^ATG6^* # 1, amiRNA*^ATG6^* # 2, *npr1*, *NPR1- GFP*, *ATG6-mCherry* and *ATG6-mCherry* × *NPR1-GFP* plants, located in the fifth and sixth positions, were infiltrated with *Pst* DC3000/*avrRps4*. After 3 days, the leaves were excised and subjected to a 1 min boiling step in trypan blue staining buffer (consisting of 10 g phenol, 10 mL glycerol, 10 mL lactic acid, 10 mL ddH_2_O, and 10 mg trypan blue), followed by destaining 3 times at 37℃ in 2.5 mg/mL chloral hydrate.

#### Statistical Analysis

All quantitative data in this study were presented as mean ± SD. The experimental data were analyzed by a two-tailed Student’s *t*-test. Significance was assigned at *P* values < 0.05 or < 0.01.

## Abbreviations

amiRNA*^ATG6^*: artificial miRNA*^ATG6^*
ATGs: Autophagy-related genes
CHX: Cycloheximide
AuNPs: Gold nanoparticles
NPR1: Nonexpressor of pathogenesis-related genes 1
NLS: Nuclear localization sequence
Pst: P. syringae pv. tomato
SINCs: SA-induced NPR1 condensates
Salicylic acid: SA
ATG6- mCherry × NPR1-GFP: UBQ10::ATG6-mCherry × 35S::NPR1-GFP

## Acknowledgments

We thank Dr. Xinnian Dong (Duke University, USA), Dr. ZhengQing Fu (University of South Carolina) and Dr. Sheng Li (South China Normal University) for their help and contribution.

## Author contribution

B.-H.Z., J.Z. and W.-L.C. designed and supervised the research; B.-H.Z. and S.-Q.H. performed majority experiments; B.-H.Z., Y.-X.M. and H.C. synthesized gold nanoparticles (AuNPs); B.-H.Z. and S.-Q.H., Y.-Z.T. performed western blotting analysis; B.-H.Z.,Y.Z and X.L performed confocal experiment; B.-H.Z. and S.-Q.H. analyzed the data; B.-H.Z. and S.-Q.H. prepared figures; B.-H.Z. and S.-Q.H. wrote the manuscript; J.Z. and W.-L.C. supervised and completed the writing; S.-Y.G. drew working model and edited language. J.Z. and W.-L.C. provided reagents/materials/analysis tools; J.Z. and W.-L.C. agrees to serve as the corresponding author responsible for contact and ensures communication.

## Data availability

The authors confirm that the data supporting the results of this study are available in the article and its supplementary materials.

## List Supplemental Information

**Supplemental Data 1**

**Supplemental Figure**

Fig. S1 Physical interaction between NPRs and ATGs in yeast.

Fig. S2 Co-localization of NPR1-GFP and ATG6-mCherry in *N.benthamiana*.

Fig. S3 The nuclear localization of ATG6 in *Arabidopsis*.

Fig. S4 Identification of *ATG6-mCherry* × *NPR1-GFP* plants.

Fig. S5 Subcellular fractionation of endogenous ATG6 in Col after 0.5 mM SA treatment for 0, 3, 6 and 20 h.

Fig S6. Confocal images of NPR1-GFP nuclear localization in 7-day-old seedlings of *NPR1-GFP* and *ATG6-mCherry* × *NPR1-GFP* under normal and 0.5 mM SA spray for 3 h.

Fig. S7 ATG6 increases the nuclear accumulation of NPR1 under SA treatment.

Fig. S8 Overexpression of *ATG6* delayed dark-induced leaf senescence. Fig. S9 Expression of *ICS1* under normal and SA treatment conditions.

Fig. S10 Expression of *PR1* and *PR5* in Col and *ATG6-mCherry* under normal and *Pst* DC3000/*avrRps4* treatment.

Fig. S11 The protein level of NPR1-GFP in *NPR1-GFP*/silencing *ATG6* and *NPR1-GFP*/Negative control.

Fig. S12 Expression of *NPR1* in Col and *ATG6-mCherry* under normal and *Pst* DC3000/*avrRps4* treatment.

Fig. S13 Partial co-localization of ATG6-mCherry and SINCs-like condensates.

Fig S14. ATG6 improves the protein stability of NPR1 in *N. benthamiana*.

Fig. S15 Structural analysis of acidic activation domains in ATG6.

Fig. S16 ATG6 and NPR1 cooperatively inhibit *Pst* DC3000/*avrRps4*-induced cell dead.

Fig. S17 NPR1-GFP degradation assay in *ATG6-mCherry* x *NPR1-GFP Arabidopsis*.

Fig. S18. Verification of ATG6 antibody specificity.

## Supplemental Table

Table S1. Plasmid in this study.

Table S2. Primers for vector construction.

Table S3. Plant Materials.

Table S4. Antibody Information.

Table S5. Primers of RT-qPCR.

## Supplemental Results

Results S1. NPR1 and its paralogues NPR3/NPR4 physically interact with multiple ATGs.

Results S2. Overexpression of *ATG6* delays carbon starvation-induced leaf senescence, and *ATG6-GFP* and *ATG6-mCherry* fusion proteins are functional.

## Supplement Methods

Methods S1. Plasmid construction

Methods S2. *Arabidopsis* screening.

Methods S3. For *Pst* DC3000/*avrRps4* culture and infiltration.

Methods S4. Yeast two-hybrid assay.

Methods S5. Prokaryotic protein expression.

Methods S6. Nuclear and cytoplasmic separation of NPR1-GFP.

Methods S7. Protein degradation analysis.

Methods S8. Protein extraction and western blotting.

Methods S9. For the treatment of 3-week-old *N. benthamiana*.

Methods S10. For the bimolecular fluorescence complementation assay.

Methods S11. For SINCs-like condensates observation.

Methods S12. For the growth of *Pst* DC3000*/avrRps4.* Methods S13. Real-Time Quantitative PCR (RT-qPCR).

**Accession numbers**

## Supplemental Moves

Supplemental Move 1. Localization of NPR1-GFP in *N. benthamiana* co-expressed NPR1-GFP and mCherry.

Supplemental Move 2. Localization of NPR1-GFP in *N. benthamiana* co-expressed NPR1-GFP and ATG6-mCherry.

## Supplemental Data 2

Vector mapping used in this study.

