## Supplemental Data 1 for "ATG6 interacting with NPR1 increases *Arabidopsis thaliana* resistance to *Pst* DC3000/*avrRps4* by increasing its nuclear accumulation and stability"

**The following supporting information is available for this article:**

**Supplemental Figure**

Figure S1. Physical interaction between NPRs and ATGs in yeast.

Figure S2. Co-localization of NPR1-GFP and ATG6-mCherry in *N.benthamiana*.

Figure S3. The nuclear localization of ATG6 in *Arabidopsis*.

Figure S4. Identification of *ATG6-mCherry* × *NPR1-GFP* plants.

Figure S5. Subcellular fractionation of endogenous ATG6 in Col after 0.5 mM SA treatment for 0, 3, 6 and 20 h.

Figure S6. Confocal images of NPR1-GFP nuclear localization in 7-day-old seedlings of *NPR1-GFP* and *ATG6-mCherry* × *NPR1-GFP* under normal and 0.5 mM SA spray for 3 h.

Figure S7. ATG6 increases the nuclear accumulation of NPR1 under SA treatment.

Figure S8. Overexpression of *ATG6* delayed dark-induced leaf senescence.

Figure S9. Expression of *ICS1* under normal and SA treatment conditions.

Figure S10. Expression of *PR1* and *PR5* in Col and *ATG6-mCherry* under normal and *Pst* DC3000/*avrRps4* treatment.

Figure S11. The protein level of NPR1-GFP in *NPR1-GFP*/silencing *ATG6* and *NPR1-GFP*/Negative control.

Figure S12. Expression of *NPR1* in Col and *ATG6-mCherry* under normal and *Pst* DC3000/*avrRps4* treatment.

Figure. S13 Partial co-localization of ATG6-mCherry and SINCs-like condensates.

Figure S14. ATG6 improves the protein stability of NPR1 in *N. benthamiana.*

Figure S15. Structural analysis of acidic activation domains in ATG6.

Figure S16. ATG6 and NPR1 cooperatively inhibit *Pst* DC3000/*avrRps4*-induced cell dead.

Figure S17. NPR1-GFP degradation assay in *ATG6-mCherry* x *NPR1-GFP* *Arabidopsis*.

Figure S18. Verification of ATG6 antibody specificity

**Supplemental Table**

Table S1. Plasmid in this study.

Table S2. Primers for vector construction.

Table S3. Plant Materials.

Table S4. Antibody Information.

Table S5. Primers of RT-qPCR.

**Supplemental Result**

Result S1. NPR1 and its paralogues NPR3/NPR4 physically interact with multiple ATGs.

Result S2. Overexpression of *ATG6* delays carbon starvation-induced leaf senescence, and *ATG6-GFP* and *ATG6-mCherry* fusion proteins are functional.

**Supplemental Methods**

Methods S1. Plasmid construction.

Methods S2. *Arabidopsis* screening.

Methods S3. For *Pst* DC3000/*avrRps4* culture and infiltration.

Methods S4. Yeast two-hybrid assay.

Methods S5. Prokaryotic protein expression.

Methods S6. Nuclear and cytoplasmic separation of NPR1-GFP.

Methods S7. Protein degradation analysis.

Methods S8. Protein extraction and western blotting.

Methods S9. For the treatment of 3-week-old *Nicotiana benthamiana*.

Methods S10. For the bimolecular fluorescence complementation assay.

Methods S11. For SINCs-like condensates observation.

Methods S12. For the growth of *Pst* DC3000*/avrRps4.*

Methods S13. Real-Time Quantitative PCR (RT-qPCR).

**Accession numbers**

**Supplemental Moves**

**Supplemental Move 1.** Localization of NPR1-GFP in *Nicotiana benthamiana* leaves co-expressed NPR1-GFP and mCherry.

**Supplemental Move 2.** Localization of NPR1-GFP in *Nicotiana benthamiana* leaves co-expressed NPR1-GFP and ATG6-mCherry.

**Supplemental Figure**

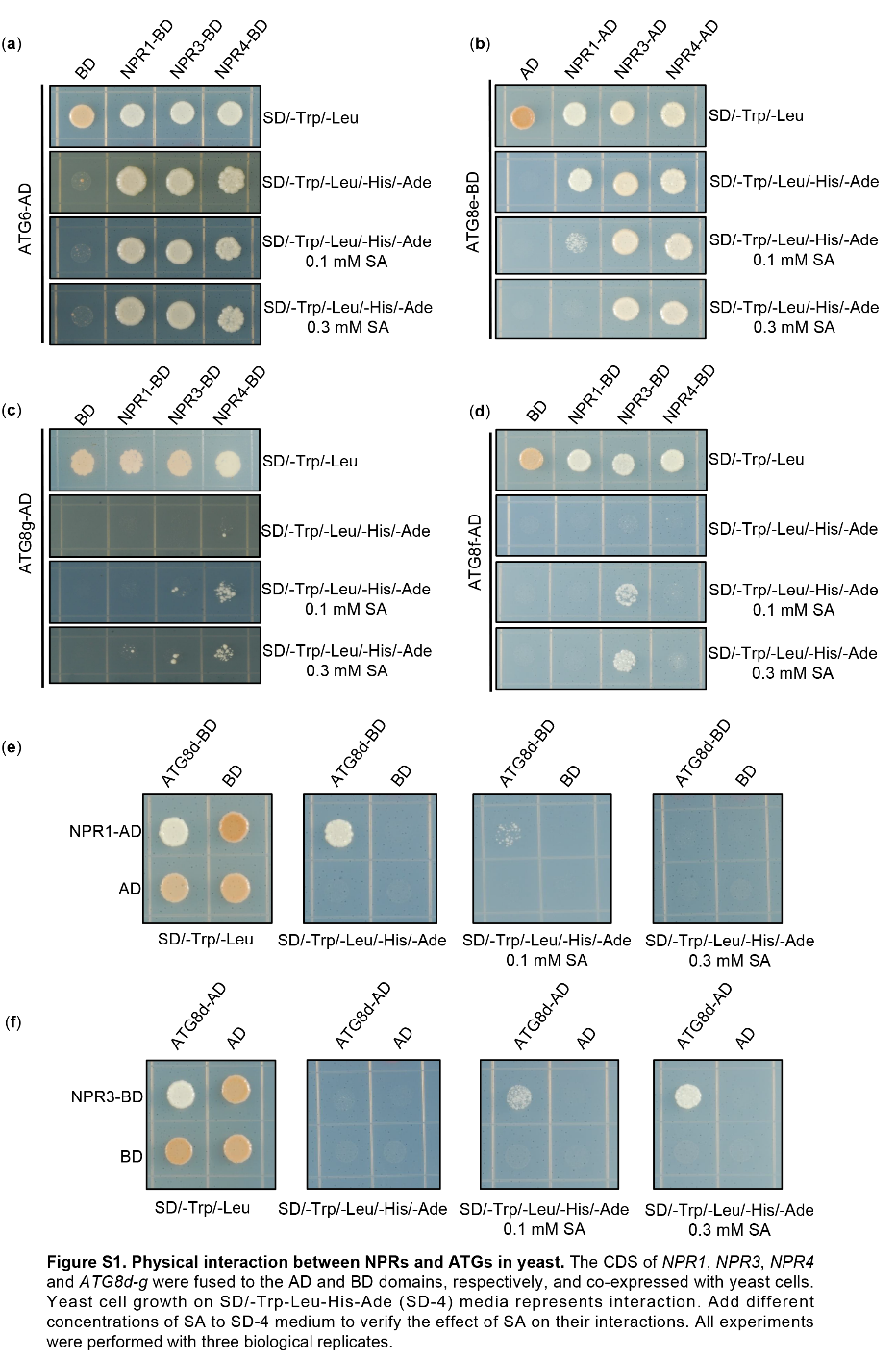

**Figure S1.** Physical interaction between NPRs and ATGs in yeast. The CDS of *NPR1*, *NPR3*, *NPR4*, *ATG6* and *ATG8d-g* were fused to the AD and BD domains, respectively, and co-expressed with yeast cells. Yeast cell growth on SD/-Trp-Leu-His-Ade media represents interaction. Add different concentrations of SA to SD/-Trp-Leu-His-Ade medium to verify the effect of SA on their interactions. All experiments were performed with three biological replicates.

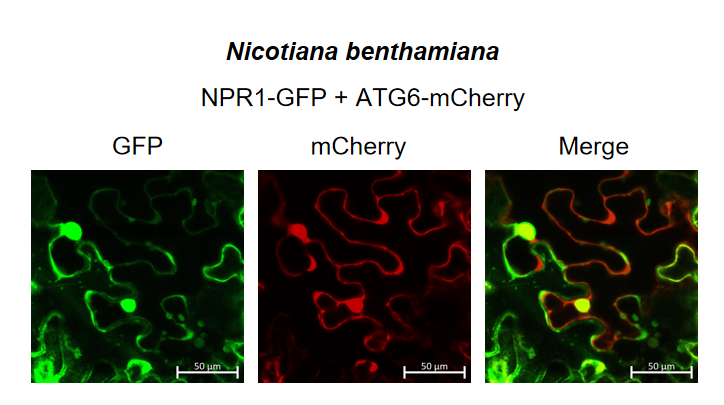

**Figure S2.** Co-localization of NPR1-GFP and ATG6-mCherry in *N.benthamiana*. NPR1-GFP were co-expressed with ATG6-mCherry in *N. benthamiana* for 3 d followed by confocal observation. Scale bar, 50 μm. Experiment was performed with three biological replicates.

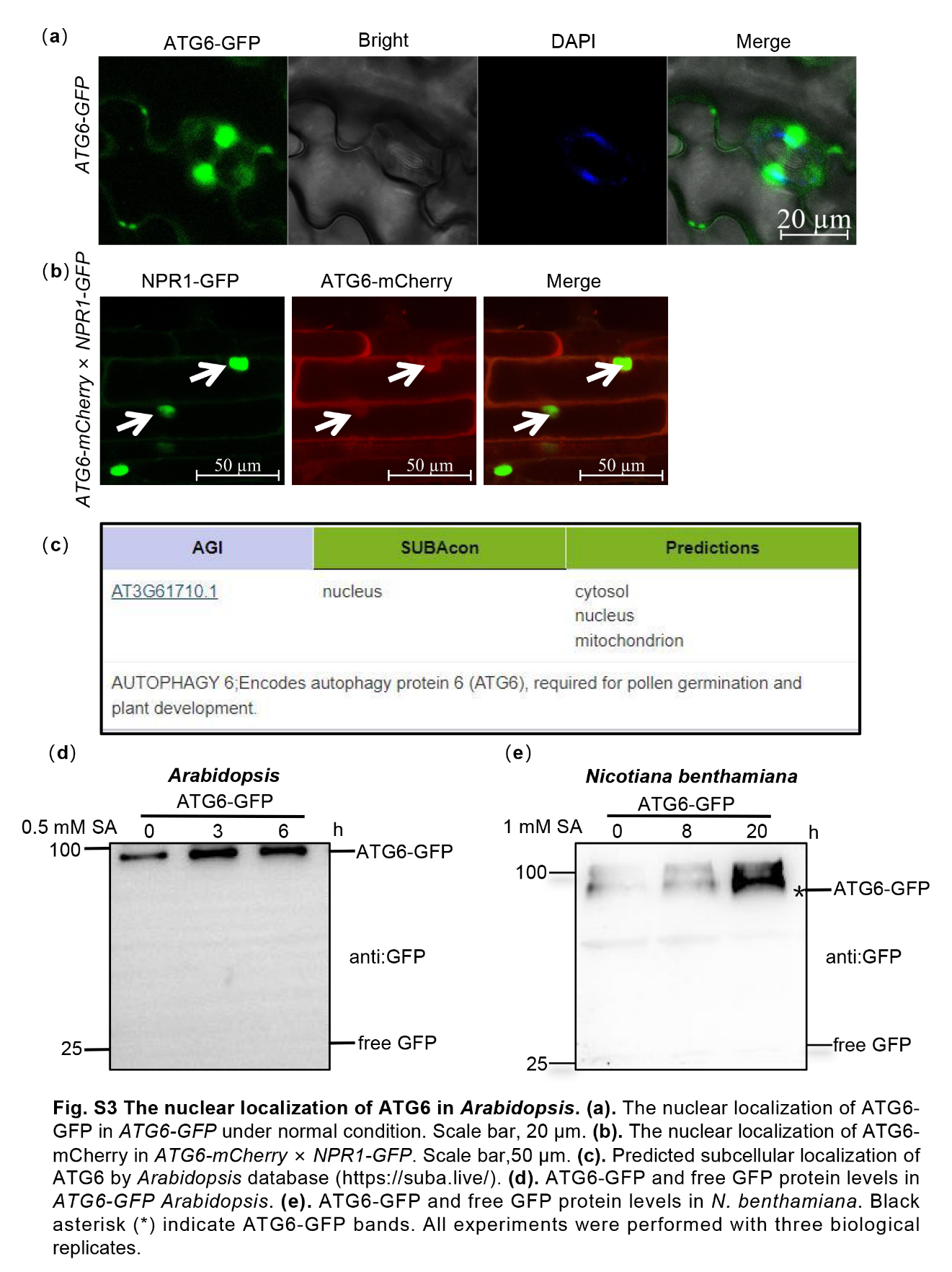

**Figure S3.** The nuclear localization of ATG6 in *Arabidopsis*. (**a**). The nuclear localization of ATG6-GFP in *ATG6-GFP* under normal condition. Scale bar, 20 μm. (**b**). The nuclear localization of ATG6-mCherry in *ATG6-mCherry* × *NPR1-GFP.* Scale bar, 50 μm. (**c**). Predicted subcellular localization of ATG6 by *Arabidopsis* database (<https://suba.live/>). (**d**). ATG6-GFP and free GFP protein levels in *ATG6-GFP* *Arabidopsis*. (**e**). ATG6-GFP and free GFP protein levels in *N. benthamiana*. Black asterisk (*) indicate ATG6-GFP bands. All experiments were performed with three biological replicates.

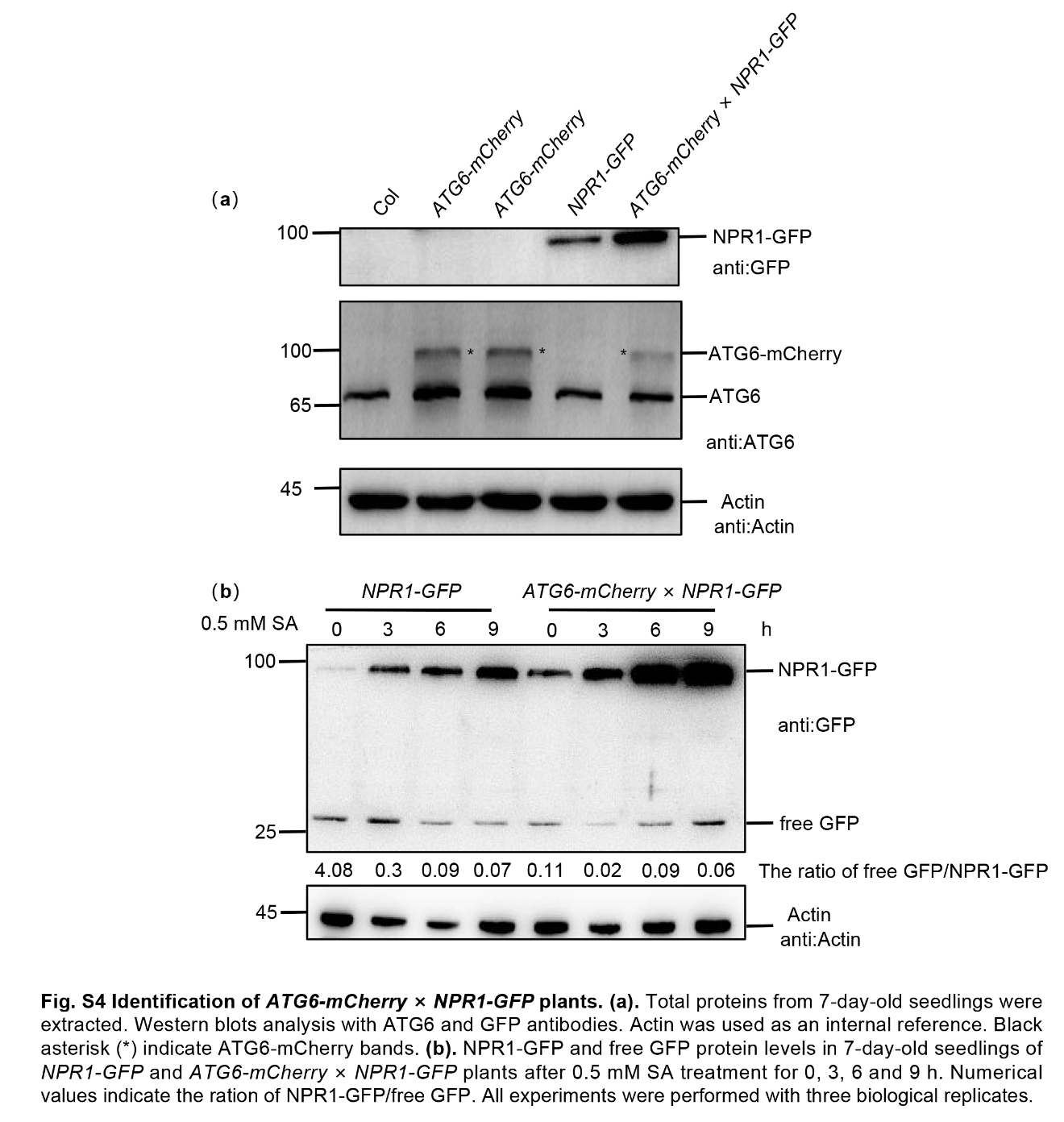

**Figure S4.** Identification of *ATG6-mCherry* × *NPR1-GFP* plants. (**a**). Total proteins from 7-day-old seedlings were extracted. Western blot was performed using ATG6 and GFP antibodies. Actin was used as an internal reference. Black asterisk (*) indicate ATG6-GFP bands. (**b**). NPR1-GFP and free GFP protein levels in 7-day-old seedlings of *NPR1-GFP* and *ATG6-mCherry* × *NPR1-GFP* plants after 0.5 mM SA treatment for 0, 3, 6 and 9 h. Numerical values indicate the ration of NPR1-GFP/free GFP. All experiments were performed with three biological replicates.

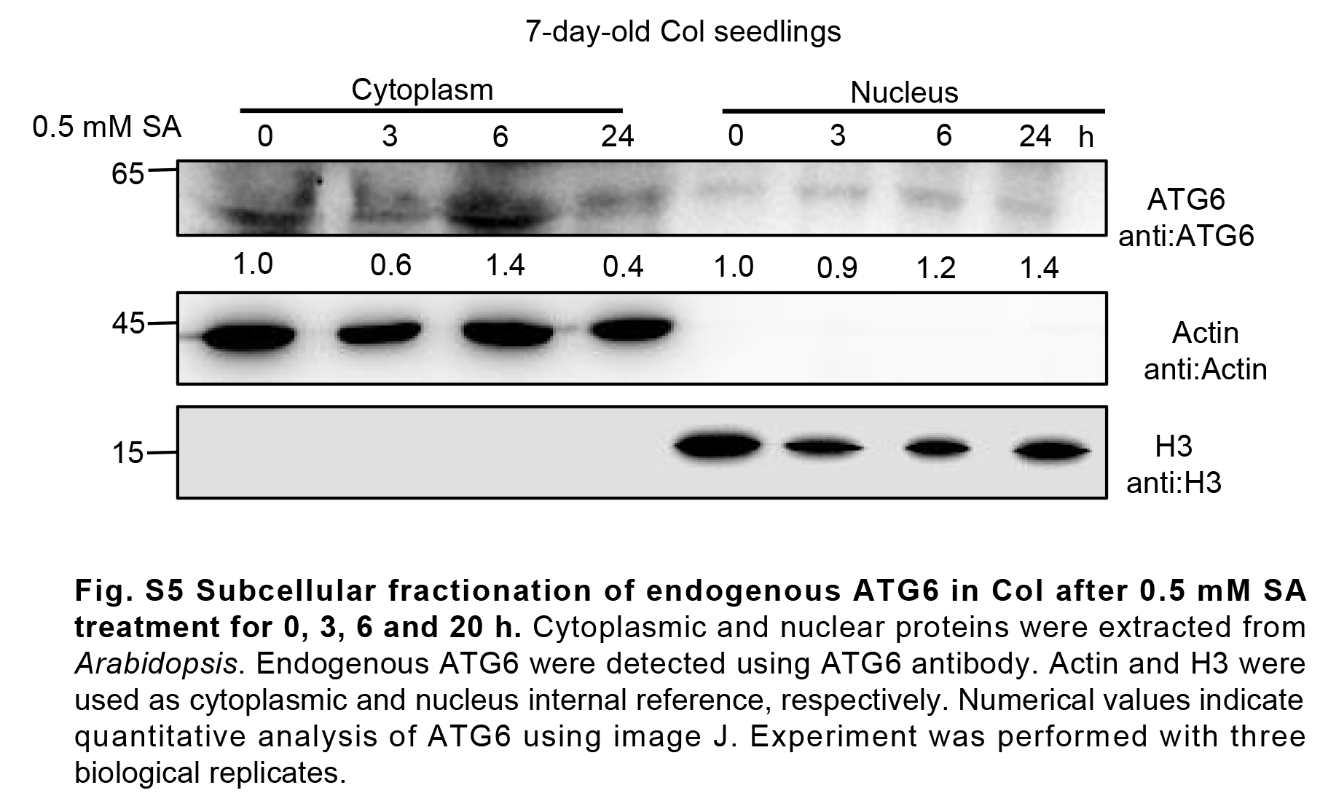

**Figure S5.** Subcellular fractionation of endogenous ATG6 in Col after 0.5 mM SA treatment for 0, 3, 6 and 20 h. Cytoplasmic and nuclear proteins were extracted from *Arabidopsis*. Endogenous ATG6 were detected using ATG6 antibody. Actin and H3 were used as cytoplasmic and nucleus internal reference, respectively. Numerical values indicate quantitative analysis of ATG6 using image J. Experiment was performed with three biological replicates.

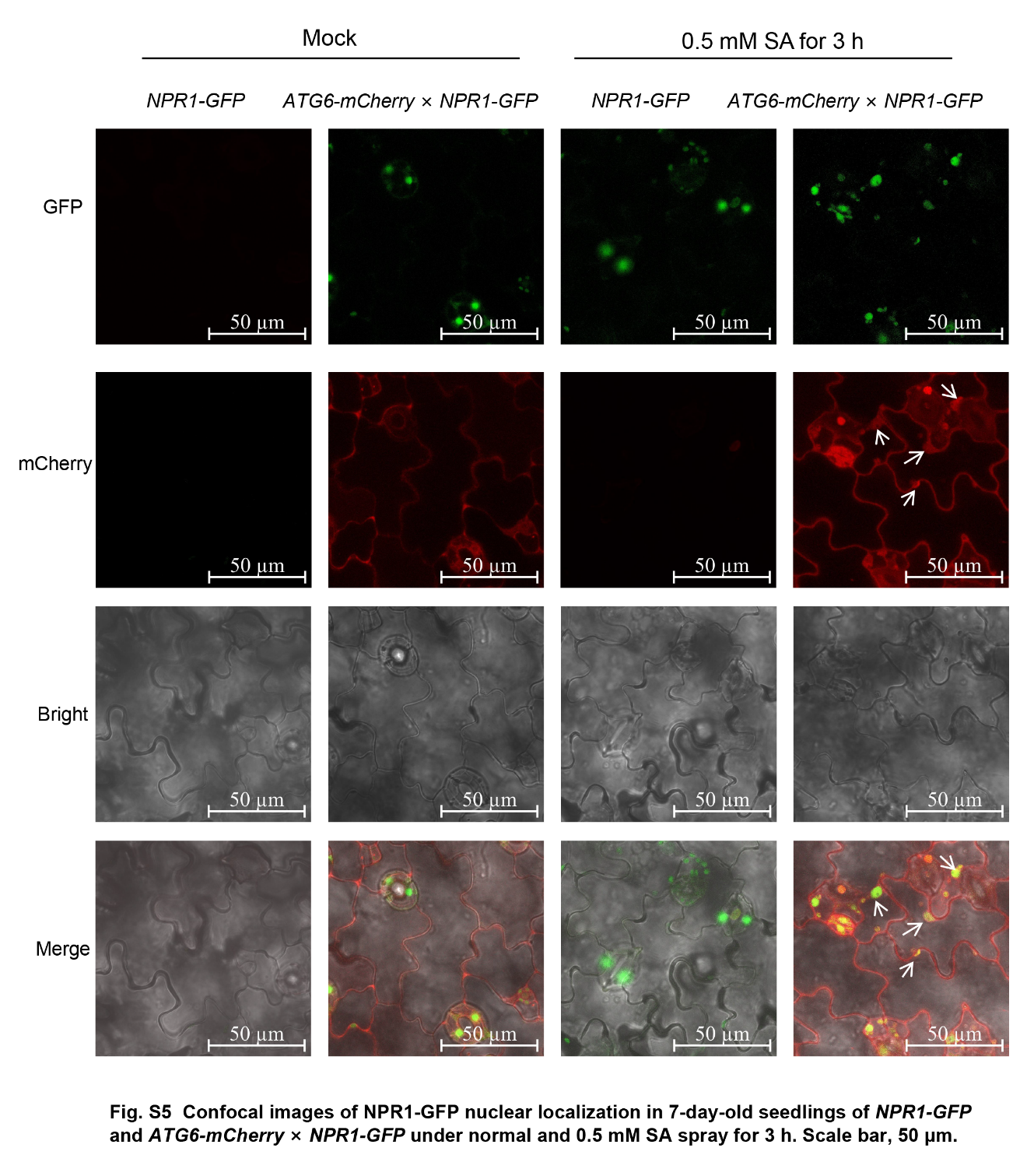

**Figure S6.** Confocal images of NPR1-GFP nuclear localization in 7-day-old seedlings of *NPR1-GFP* and *ATG6-mCherry* × *NPR1-GFP* under normal and 0.5 mM SA spray for 3 h. Scale bar, 50 μm. Experiment was performed with three biological replicates.

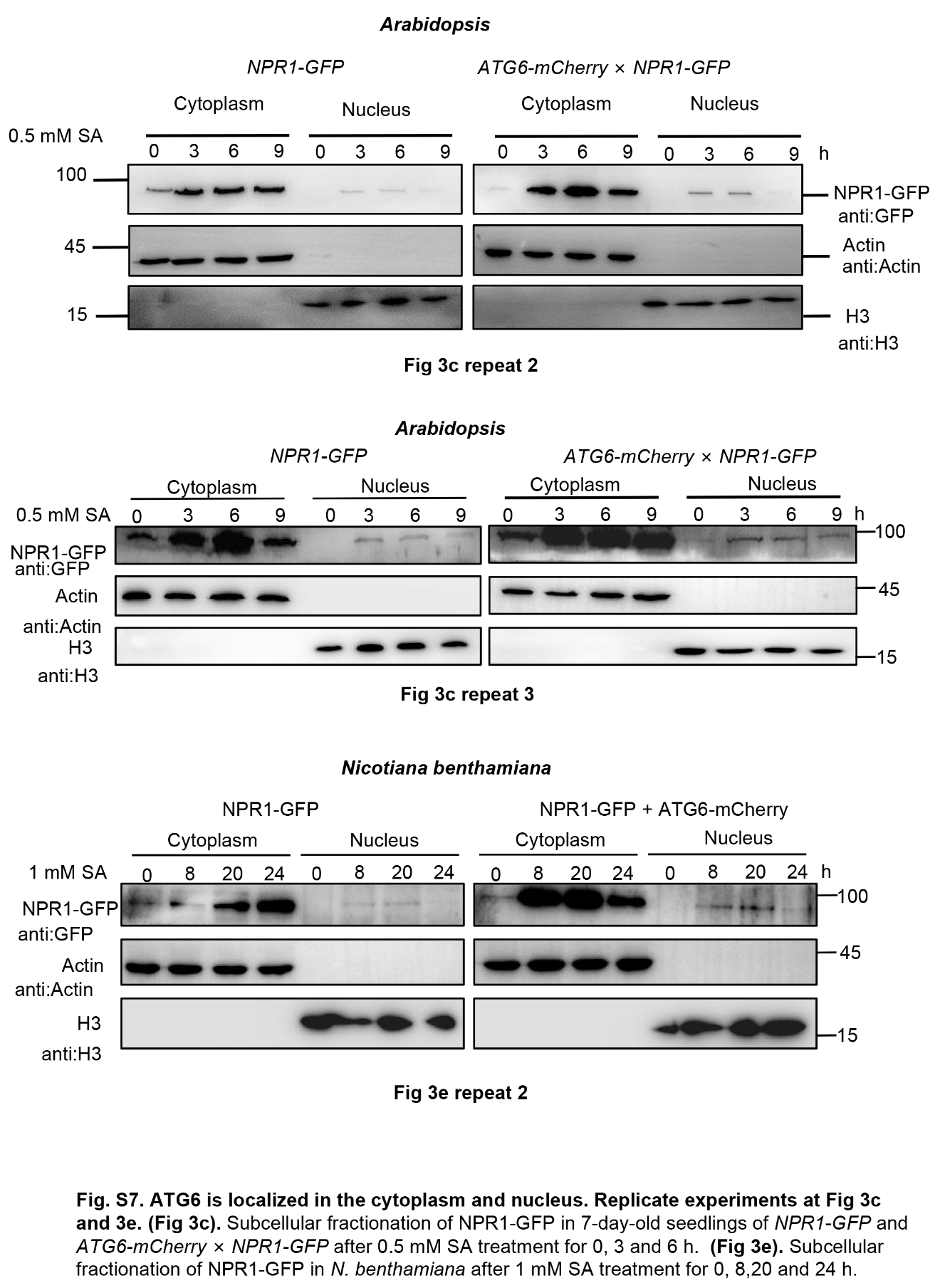

**Figure S7**. ATG6 increases the nuclear accumulation of NPR1 under SA treatment. Replicate experiments at Fig 3c and 3e. Fig 3c. Subcellular fractionation of NPR1-GFP in 7-day-old seedlings of *NPR1-GFP* and *ATG6-mCherry* × *NPR1-GFP* after 0.5 mM SA treatment for 0, 3 and 6 h. Fig 3e Subcellular fractionation of NPR1-GFP in *N. benthamiana* after 1 mM SA treatment for 0, 8,20 and 24 h.

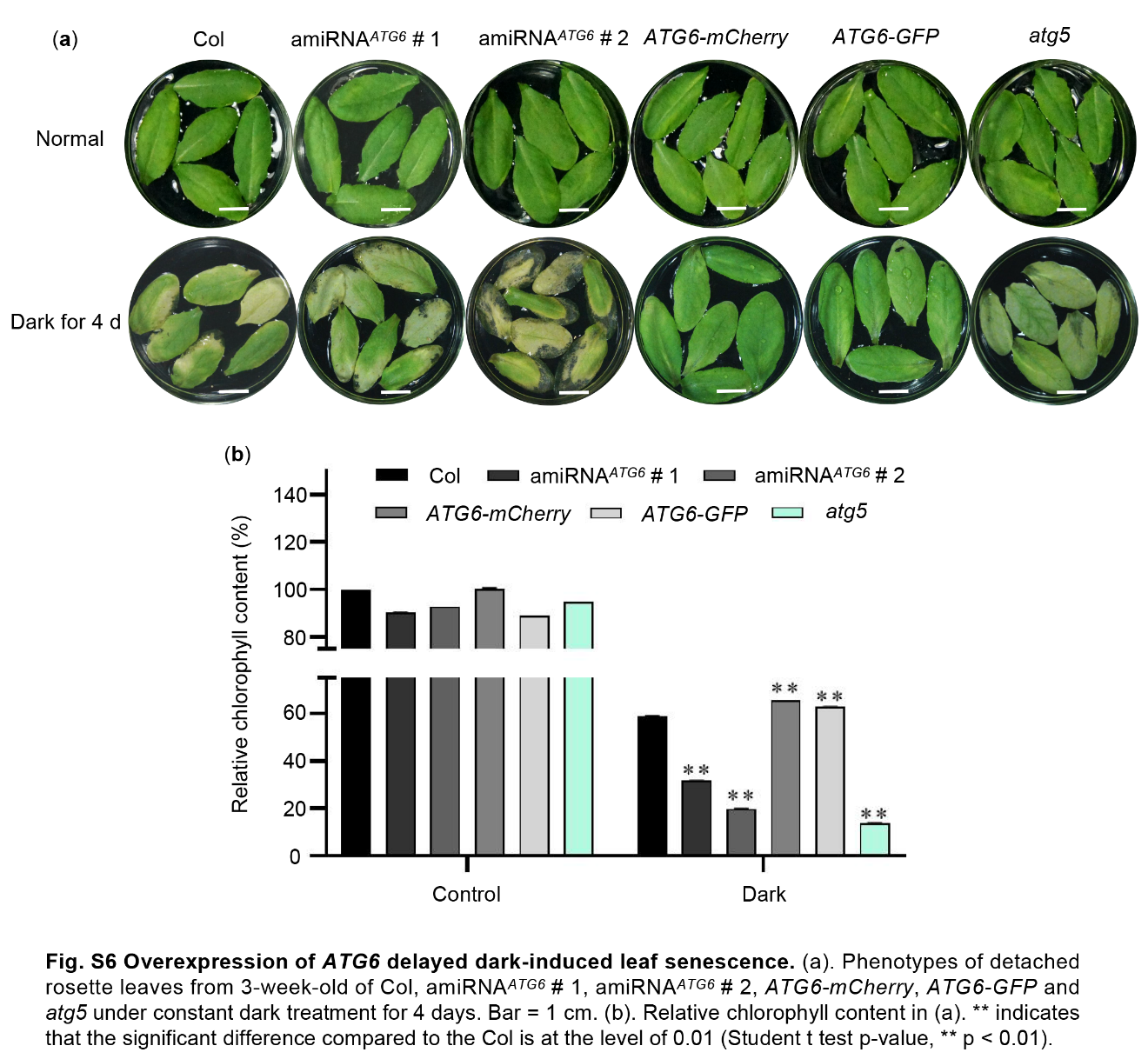

**Figure S8.** Overexpression of *ATG6* delayed dark-induced leaf senescence. (a). Phenotypes of detached rosette leaves from 3-week-old of Col, amiRNA*^ATG6^* # 1, amiRNA*^ATG6^* # 2, *ATG6-mCherry*, *ATG6-GFP* and *atg5* under constant dark treatment for 4 days. Bar = 1 cm. (b). Relative chlorophyll content in (a). ** indicates that the significant difference compared to the Col is at the level of 0.01 (Student *t* test *p*-value, ** *p* < 0.01). Experiment was performed with three biological replicates.

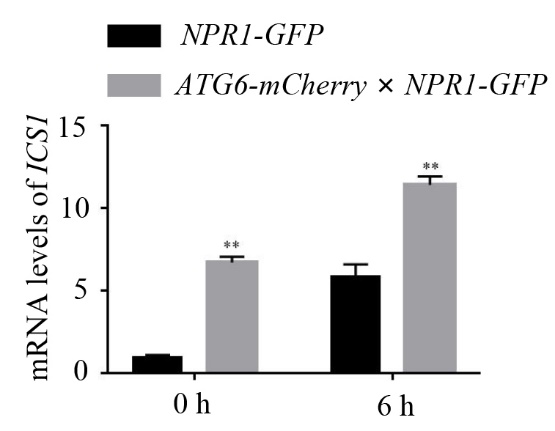

**Figure S9.** Expression of *ICS1* in 3-week-old *NPR1-GFP* and *ATG6-mCherry* × *NPR1-GFP* under normal and 0.5 mM SA treatment conditions. Values are means ± SD (n = 3 biological replicates). The *AtActin* gene was used as the internal control. ** indicates that the significant difference compared to the control is at the level of 0.01 (Student *t* test *p* value, ** *p*< 0.01). Experiment was performed with three biological replicates.

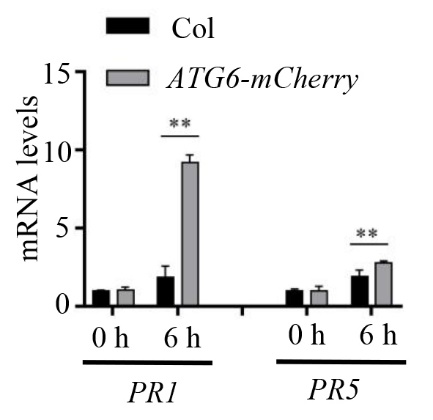

**Figure S10.** Expression of *PR1* and *PR5* in Col and *ATG6-mCherry* under normal and *Pst* DC3000/*avrRps4* treatment. Values are means ± SD (n = 3 biological replicates). The *AtActin* gene was used as the internal control. ** indicates that the significant difference compared to the control is at the level of 0.01 (Student *t* test *p* value, ** *p*< 0.01). Experiment was performed with three biological replicates.

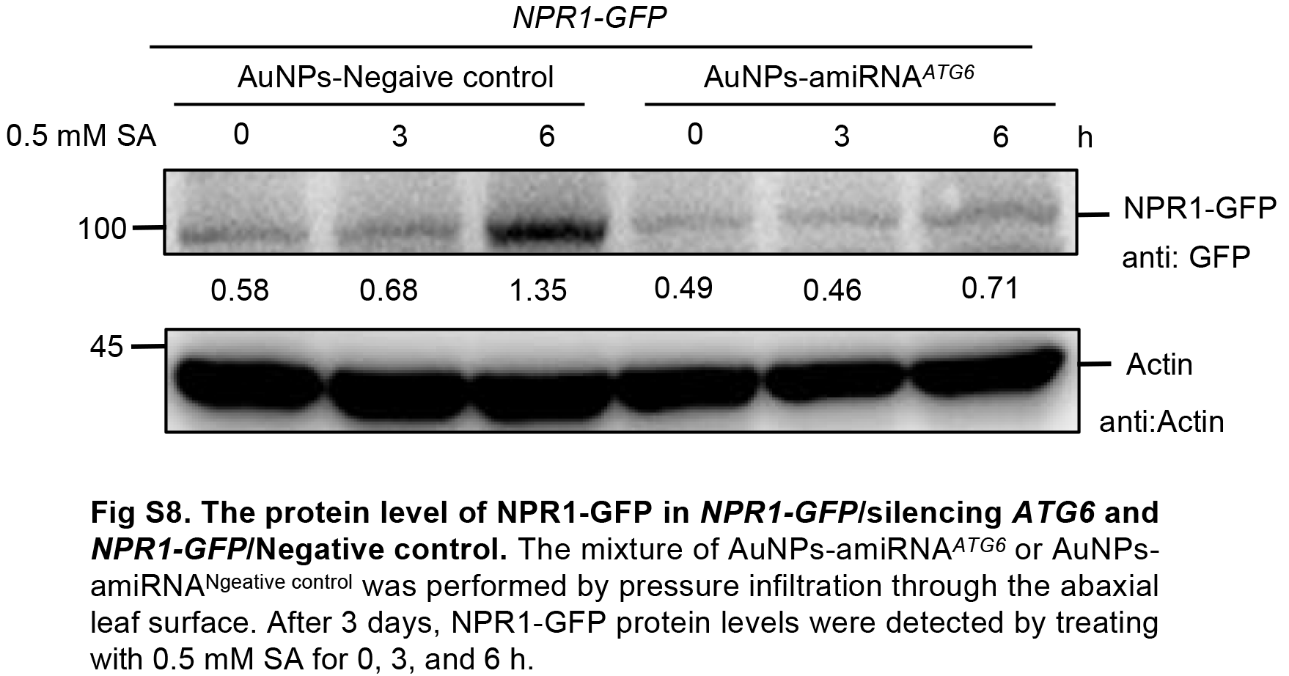

**Figure S11.** The protein level of NPR1-GFP in *NPR1-GFP*/silencing *ATG6* and *NPR1-GFP*/Negative control. The mixture of AuNPs-amiRNA*^ATG6^* or AuNPs-amiRNA^Ngeative control^ was performed by pressure infiltration through the abaxial leaf surface. After 3 days, NPR1-GFP protein levels were detected by treating with 0.5 mM SA for 0, 3, and 6 h. Experiment was performed with three biological replicates.

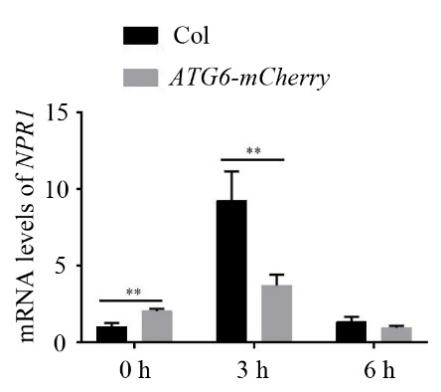

**Figure S12.** Expression of *NPR1* in Col and *ATG6-mCherry* under normal and *Pst* DC3000/*avrRps4* treatment. Values are means ± SD (n = 3 biological replicates). The *AtActin* gene was used as the internal control. ** indicates that the significant difference compared to the control is at the level of 0.01 (Student *t* test *p* value, ** *p*< 0.01). Experiment was performed with three biological replicates.

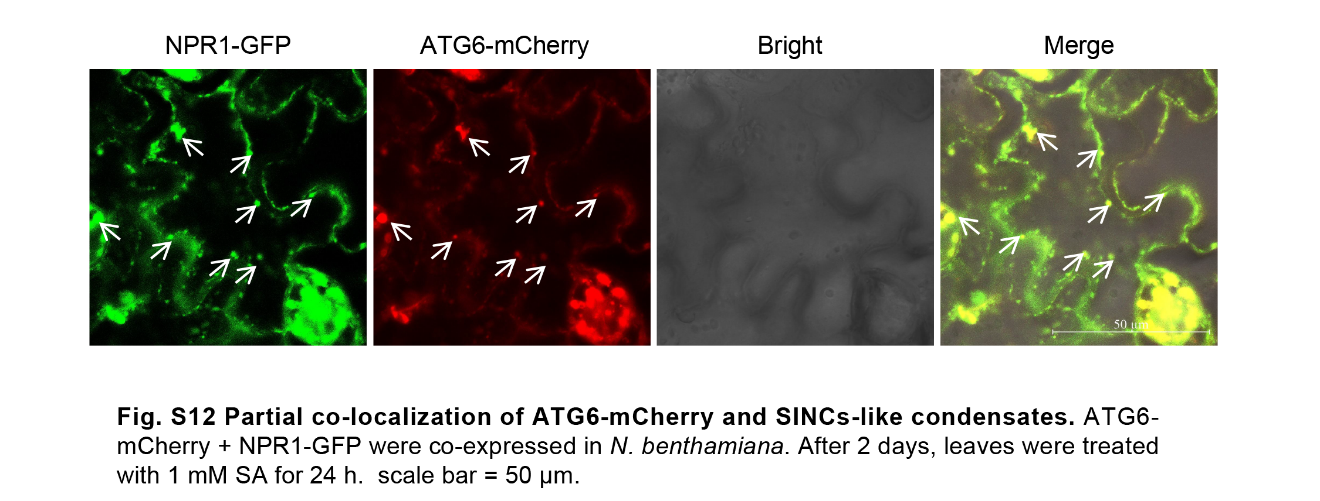

**Figure S13.** Partial co-localization of ATG6-mCherry and SINCs-like condensates. ATG6-mCherry + NPR1-GFP were co-expressed in *N. benthamiana*. After 2 days, leaves were treated with 1 mM SA for 24 h. scale bar = 50 μm. Experiment was performed with three biological replicates.

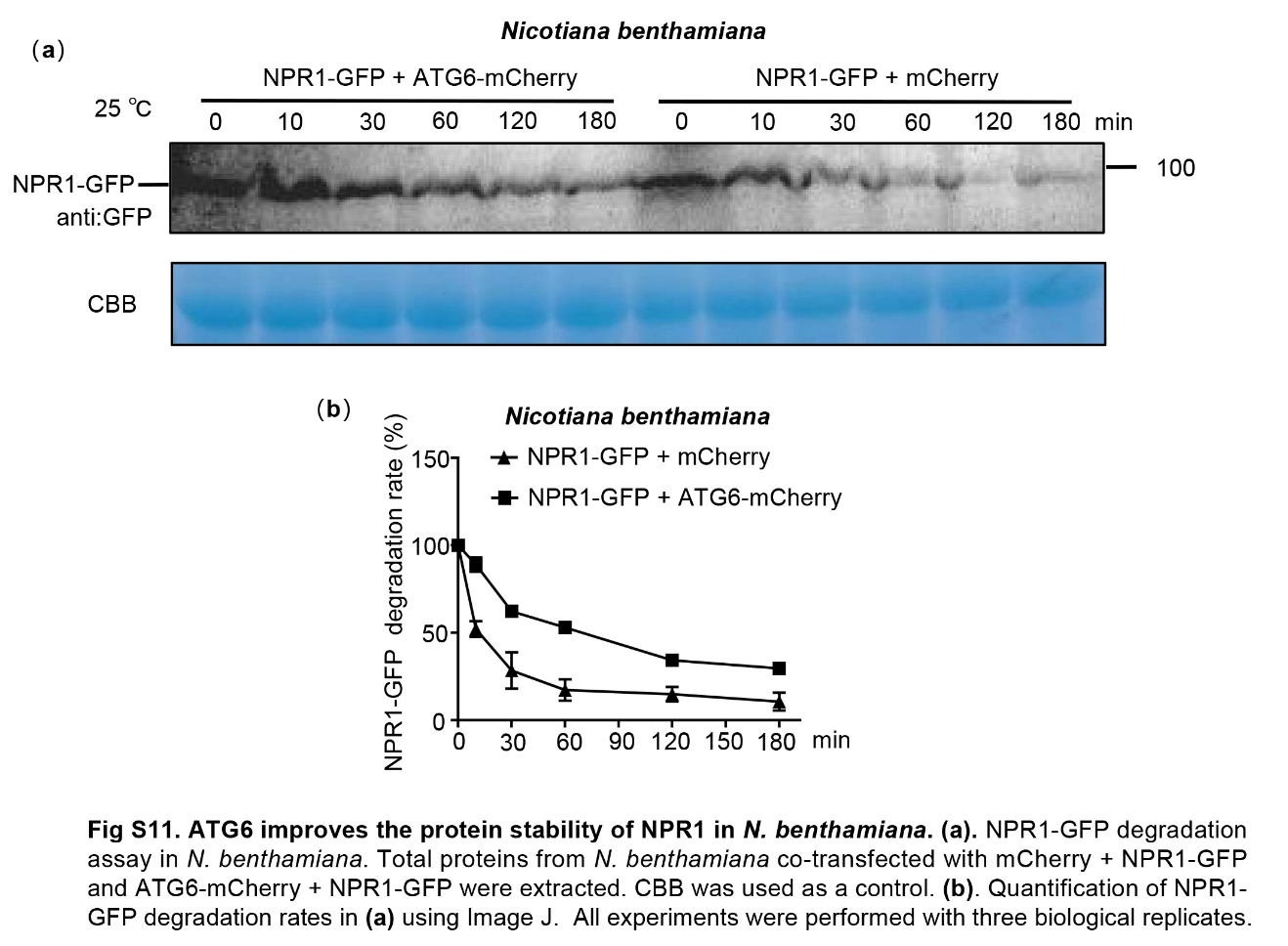

**Figure S14.** ATG6 improves the protein stability of NPR1 in *N. benthamiana*. (a). NPR1-GFP degradation assay in *N. benthamiana*. Total proteins from *N. benthamiana* co-transfected with mCherry + NPR1-GFP and ATG6-mCherry + NPR1-GFP were extracted. CBB was used as a control. (b). Quantification of NPR1-GFP degradation rates in (a) using Image J. All experiments were performed with three biological replicates.

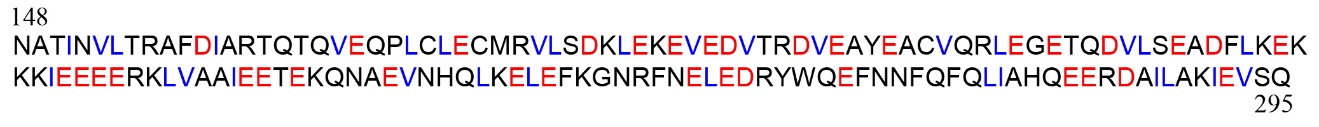

**Figure S15.** Structural analysis of acidic activation domains (AADs) in ATG6. Acidic (red) and hydrophobic (blue) amino acid residues in AADs.

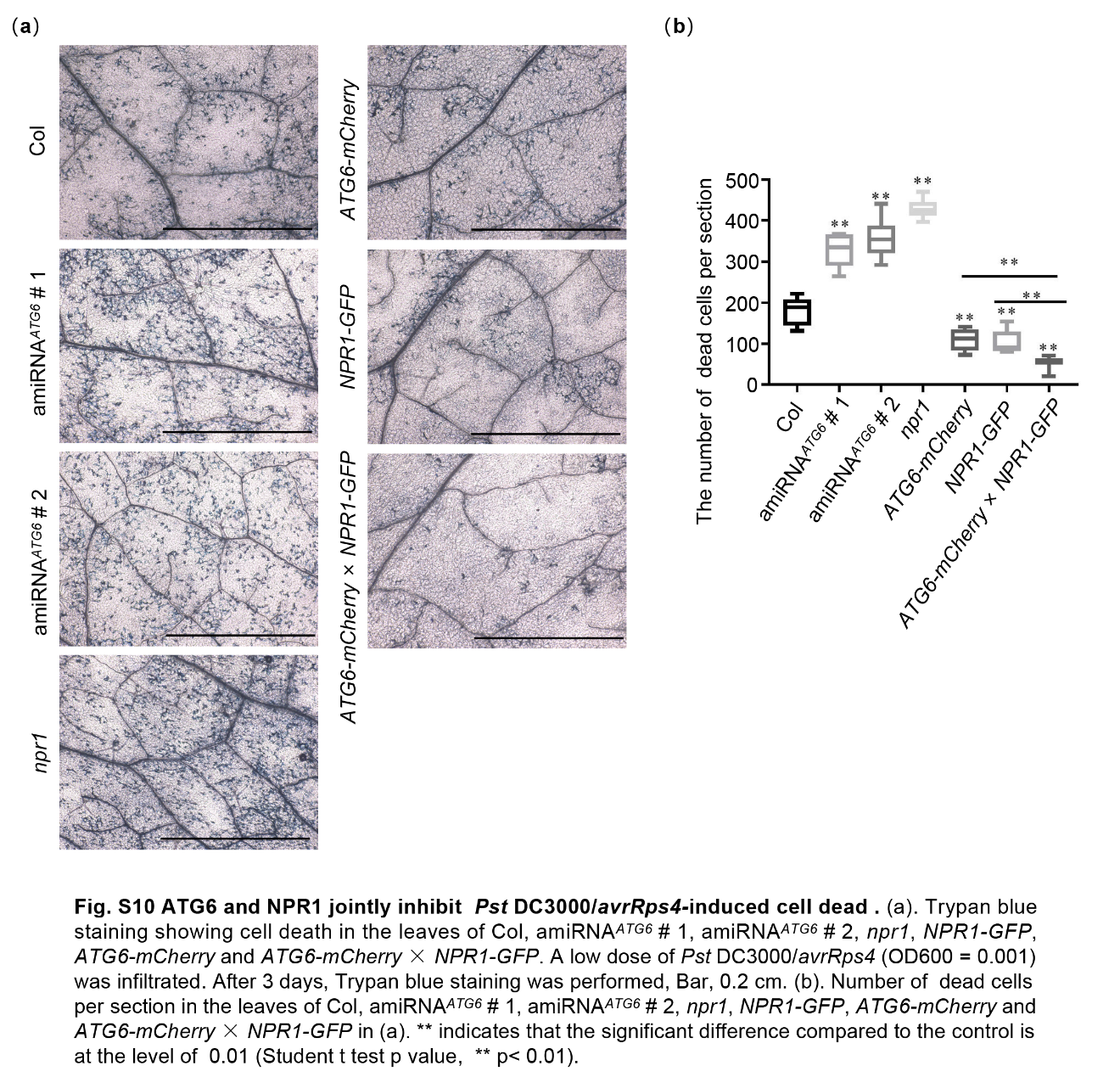

**Figure S16.** ATG6 and NPR1 cooperatively inhibit *Pst* DC3000/*avrRps4*-induced cell dead. (a). Trypan blue staining showing cell death in the leaves of Col, amiRNA*^ATG6^* # 1, amiRNA*^ATG6^* # 2, *npr1*, *NPR1-GFP*, *ATG6-mCherry* and *ATG6-mCherry* × *NPR1-GFP*. A low dose of *Pst* DC3000/*avrRps4* (OD600 = 0.001) was infiltrated. After 3 days, Trypan blue staining was performed, Bar, 0.2 cm. (b). Number of dead cells per section in the leaves of Col, amiRNA*^ATG6^* # 1, amiRNA*^ATG6^* # 2, *npr1*, *NPR1-GFP*, *ATG6-mCherry* and *ATG6-mCherry* × *NPR1-GFP* in (a). ** indicates that the significant difference compared to the control is at the level of 0.01 (Student t test p value, ** *p< 0.01*). Experiment was performed with three biological replicates.

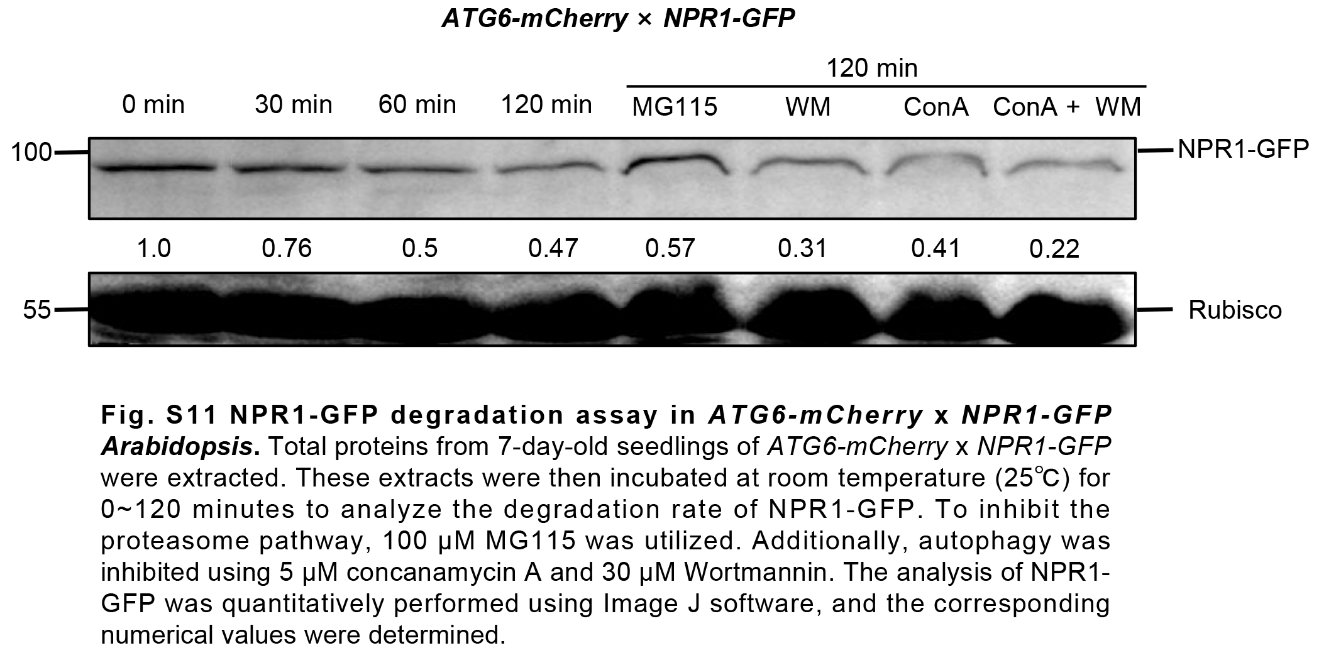

**Figure S17.** NPR1-GFP degradation assay in *ATG6-mCherry* x *NPR1-GFP* *Arabidopsis*. Total proteins from 7-day-old seedlings of *ATG6-mCherry* x *NPR1-GFP* were extracted. These extracts were then incubated at room temperature (25℃) for 0~120 minutes to analyze the degradation rate of NPR1-GFP. To inhibit the proteasome pathway, 100 μM MG115 was utilized. Additionally, autophagy was inhibited using 5 μM concanamycin A and 30 μM Wortmannin. The analysis of NPR1-GFP was quantitatively performed using Image J software, and the corresponding numerical values were determined. Experiment was performed with three biological replicates.

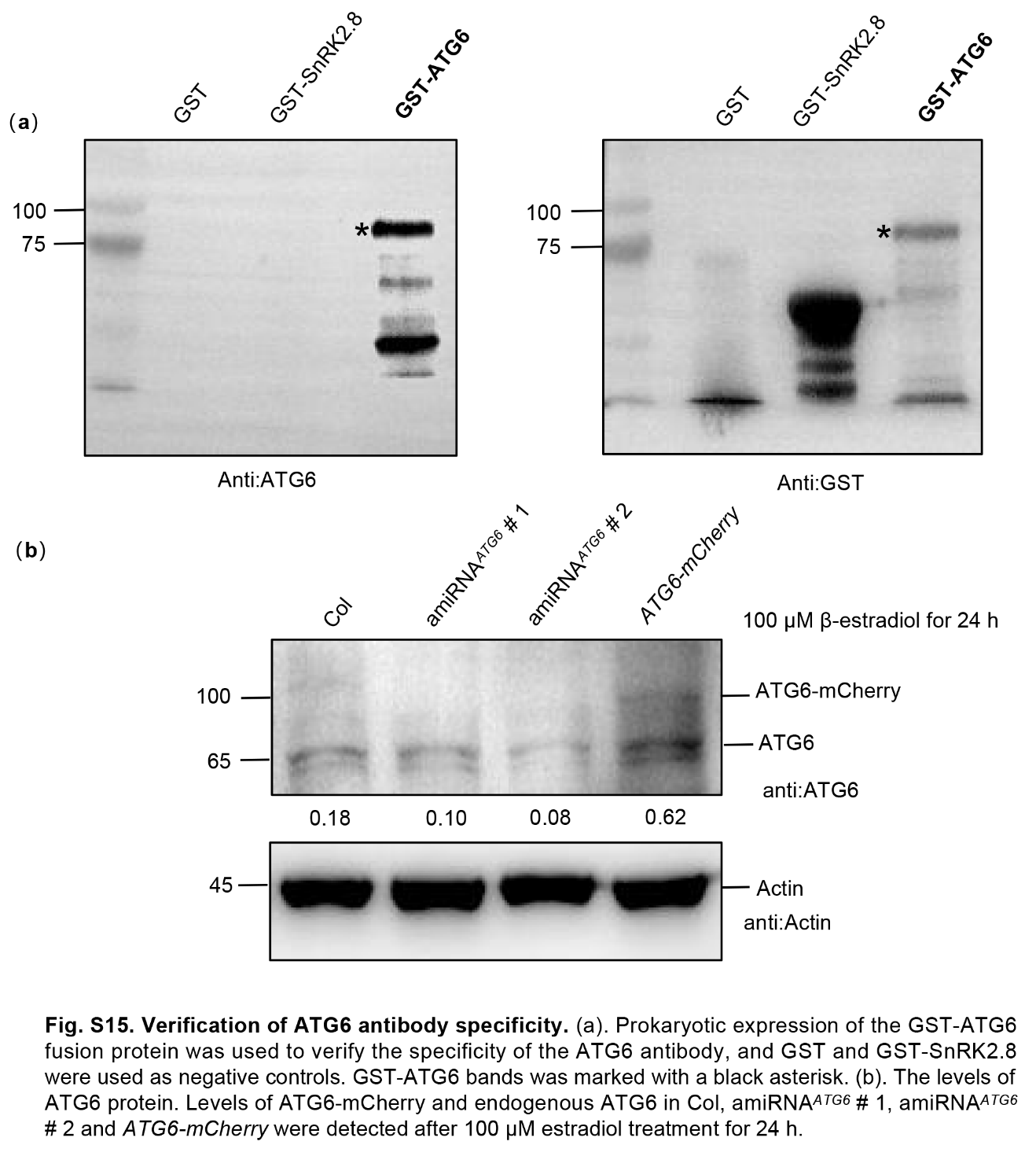

**Figure S18**. Verification of ATG6 antibody specificity. (a). Prokaryotic expression of the GST-ATG6 fusion protein was used to verify the specificity of the ATG6 antibody, and GST and GST-SnRK2.8 were used as negative controls. GST-ATG6 bands was marked with a black asterisk. (b). The levels of ATG6 protein. Levels of ATG6-mCherry and endogenous ATG6 in Col, amiRNAATG6 # 1, amiRNAATG6 # 2 and ATG6-mCherry were detected after 100 μM estradiol treatment for 24 h. Experiment was performed with three biological replicates.

**Supplemental Table**

**Table S1**. Plasmid in this study.

| Gene name | | Vector Name | Subcloning method | Source |
| --- | --- | --- | --- | --- |
| *ATG6* | | pGADT7 | ClonExpress II One Step Cloning Kit | This paper |
| *NPR1* | | pGBKT7 | ClonExpress II One Step Cloning Kit | This paper |
| *NPR1-C* | | pGBKT7 | ClonExpress II One Step Cloning Kit | This paper |
| *NPR1-N* | | pGBKT7 | ClonExpress II One Step Cloning Kit | This paper |
| *SnRK2.8* | | pGADT7 | ClonExpress II One Step Cloning Kit | This paper |
| *ATG6* | | 35s-gene-cYFP | ClonExpress II One Step Cloning Kit | This paper |
| *NPR1* | | 35s-gene-nYFP | ClonExpress II One Step Cloning Kit | This paper |
| *SnRK2.8* | | 35s-gene-cYFP | ClonExpress II One Step Cloning Kit | This paper |
| *SnRK2.8* | | pGEX-4T-1 | Double digests and T4 DNA ligase | This paper |
| *ATG6* | | pGEX-4T-1 | Double digests and T4 DNA ligase | This paper |
| *ATG6* | 1300:UBQ-mCherry | | Double digests and T4 DNA ligase | This paper |
| *ATG6* | | 1300:UBQ-eGFP | Double digests and T4 DNA ligase | This paper |
| *GFP* | | 1300:UBQ-eGFP | N/A | This paper |
| *NPR1* | | pET32a | Provided by Dr. ZhengQing Fu of University of South Carolina | |
| *NPR1*  amiRNA*^ATG6^* | | pCB302-GFP  pERM10M | Provided by Dr. ZhengQing Fu of University of South Carolina  Double digests and T4 DNA ligase This paper | |

**Table S2.** Primers for vector construction.

| Vector Name | Primers（5′-3′） |
| --- | --- |
| For Y2H assay | |
| ATG6-AD (pGADT7） | F: GAGGCCAGTGAATTCCACCCGATGAGGAAAGAGGAGATTCCAG  R: CCCGTATCGATGCCCACCCCTAAGTTTTTTTACATGAAGGCT |
| NPR1-BD (pGBKT7) | F: CATGGAGGCCGAATTCCCGATGGACACCACCATTGATG  R: CAGGTCGACGGATCCCCTCACCGACGACGATGAGAG |
| NPR1-C-BD (pGBKT7) | F: CATGGAGGCCGAATTCCCGCTTCATTTCGCTGTTGCAT |
| NPR1-N-BD (pGBKT7) | R: CAGGTCGACGGATCCCCTCAAGCACACGCATCATCTAGAT |
| SnRK2.8-AD (pGADT7） | F: CATGGAGGCCGAATTCCCGATGGAGAGGTACGAAATAGTGAAG  R: CAGGTCGACGGATCCCCTCACAAAGGGGAAAGGAGATCAGCGGT |
| For BiFC assay | |
| ATG6 -cYFP  (35s-gene-cYFP) | F: CGACGGTACCGCGGGCCCGGGATGAGGAAAGAGGAGATTCCAG  R: CACGCTGCCCAGGATCCCGGGAGTTTTTTTACATGAAGGCT |
| NPR1-nYFP (35s-gene-nYFP) | F: CGACGGTACCGCGGGCCCGGGATGGACACCACCATTGATG  R: GCTCACCATCAGGATCCCGGGCCGACGACGATGAGAGAG |
| SnRK2.8 -cYFP  (35s-gene-cYFP) | F: CGACGGTACCGCGGGCCCGGGATGGAGAGGTACGAAATAGTGAAG  R: CACGCTGCCCAGGATCCCGGGCAAAGGGGAAAGGAGATCAGCGGT |
| For plant transformation | |
| ATG6-mCherry  (1300:UBQ-mCherry) ATG6-GFP  (1300:UBQ-GFP)  amiRNA*^ATG6^* I  amiRNA*^ATG6^* II  amiRNA*^ATG6^* III  amiRNA^ATG6^ IV  miRNA 319 F  miRNA 319 R | F: cagACTAGTATGAGGAAAGAGGAGATTCCAG  R: cagACTAGTAGTTTTTTTACATGAAGGCTTACTAG  F: cagACTAGTATGAGGAAAGAGGAGATTCCAG  R: cagACTAGTAGTTTTTTTACATGAAGGCTTACTAG  miR-s: GATCAATTCTAGGATAACTGCCCCTCTCTTTTGTATTCCA  miR-a: AGGGGCAGTTATCCTAGAATTGATCAAAGAGAATCAATGA  miR*s: AGGGACAGTTATCCTTGAATTGTTCACAGGTCGTGATATG  miR*a: GAACAATTCAAGGATAACTGTCCCTACATATATATTCCTA  F: CGCGGATCCCAAACACACGCTCGGACGCATATT  R: TCCCCCGGGCATGGCGATGCCTTAAATAAAGATAAACCC |
| GST-ATG6 (pGEX-4T-1) | F: GAATTCATGAGGAAAGAGGAGATTCC  R: GTCGACAGTTTTTTTACATGAAGGCTTACTAG |
| GST-SnRK2.8 (pGEX-4T-1) | F: CGCGGATCCATGGAGAGGTACGAAATAGTGAAG  R: CCGCTCGAGCAAAGGGGAAAGGAGATCAGCGGT |

**Table S3.** Plant Materials.

| Name | Source |
| --- | --- |
| *NPR1-GFP/atg5* | This paper (crossing) |
| *ATG6-mCherry × NPR1-GFP/npr1-2* | This paper (crossing) |
| *ATG6-mCherry*  *ATG6-GFP*  amiRNA*^ATG6^* | This paper (floral dip method)  This paper (floral dip method)  This paper (floral dip method) |
| *atg5-1* | SALK_020601C |
| *NPR1-GFP* (in *npr1-2* background) | Provided By Dr. Xinnian Dong of Duke University |
| *npr1-1* | Provided By Dr. Xinnian Dong of Duke University |

**Table S4.** Antibody Information.

| **Antibodies** | **Dilution** | **Identifier** | **Source** |
| --- | --- | --- | --- |
| anti-GFP | 1:3000 | CAT#A-6455 | Invitrogen |
| anti-GST | 1:5000 | CAT#AT0027 | Engibody |
| anti-His | 1:2000 | CAT#AH367 | Beyotime |
| anti-Actin | 1:3000 | CAT#AT0004 | Engibody |
| anti-H3 | 1:3000 | CAT#NB500-171 | Novus Biologicals |
| anti-ATG6 | 1:200 | peptide, C-KEKKKIEEEERK | Abmart |

**Table S5.** Primers of RT-qPCR.

| Genes | Primers（5′-3′） |
| --- | --- |
| *AtActin2*  *AtNPR1* | F: GGTAACATTGTGCTCAGTGGTGG  R: AACGACCTTAATCTTCATGCTGC  F:GATCGCAAAACAAGCCACTATGG  R:ATCGAGCAGCGTCATCTTCAATT |
| *AtATG6* | F:TCCTCCATACGATGTGTAACTATTTCC  R:GCTCATAAGTTTCGTTGTTGCTGT |
| *AtPR1*  *AtPR5*  *AtICS1* | F:TGTAGCTCTTGTAGGTGCTC  R:AACTCCATTGCACGTGTTCG  F:AGTTCCTCCCGTCACTCTGG  R:TCCTCCGGATGGTCTTATCC  F: GAGACTTACGAAGGAAGATGATGAG  R:TGATCCCGACTGCAAATTCACTCTC |

**Supplemental Results**

**Result S1. NPR1** **and** **its paralogues NPR3/NPR4 physically interact with multiple ATGs**

NPR1 and its paralogues NPR3/NPR4, which frequently interact with other proteins to regulate plant immune responses (Backer *et al.*, 2019; Chen *et al.*, 2019). To identify ATGs that interact with NPRs, we performed yeast two-hybrid (Y2H) screens using NPRs as bait. Interestingly, ATG6 interacted with NPR1, NPR3 and NPR4, respectively, and different concentrations of SA treatment did not significantly affect their interaction (**Fig. S1a**). ATG8e interacted with NPR3 and NPR4, respectively, and there was no significant effect of different concentrations of SA treatment on their interactions (**Fig. S1b**). NPR1 interacted with ATG8e and ATG8d, respectively, and their interactions were inhibited with increasing SA content (**Fig. S1b and e**). NPR3 did not interact with ATG8f, ATG8d and ATG8g under normal conditions, but SA significantly promoted their interactions (**Fig. S1c-f**). ATG8g had a weak interaction with NPR4 under normal conditions, and SA significantly promoted their interactions (**Fig. S1c**). NPR1 did not interact with ATG8g and ATG8f under normal and SA treatment (**Fig. S1c and d**). NPR4 did not interact with ATG8f under normal and SA treatment (**Fig. S1d**). NPR1 is an important positive regulator of the plant immune response (Chen *et al.*, 2021b). So far, nine ATG8 isoforms have been identified in *Arabidopsis*, and considering the possible redundancy between ATG8 protein (Bu *et al.*, 2020), we mainly further investigated the function of ATG6 interactions with NPR1 in plant immune response.

**Result S2. Overexpression of *ATG6* delays carbon starvation-induced leaf senescence, and *ATG6-GFP* and *ATG6-mCherry* fusion proteins are functional.**

AtATG6 is a member of the class III phosphatidylinositol 3-kinase family (PtdIns3K), which regulates autophagosome nucleation in *Arabidopsis* (Qi et al., 2017; Bozhkov, 2018). Previous studies have shown that one of the most prominent features of autophagy-deficient mutants is hypersensitivity to carbon starvation with premature senescence, and shorter growth cycles (Yoshimoto *et al.*, 2009; Bozhkov, 2018; Huang *et al.*, 2019). In contrast, activated autophagy delays carbon starvation-induced leaf senescence (Yoshimoto *et al.*, 2009; Bozhkov, 2018; Huang *et al.*, 2019). To verify whether the ATG6-GFP and ATG6-mCherry fusion proteins are functional in *Arabidopsis*. We analyzed phenotypic changes of Col, amiRNA*^ATG6^* # 1, amiRNA*^ATG6^* # 2, *ATG6-GFP* and *ATG6-mCherry* under carbon starvation. Autophagy-deficient mutant *atg5* was used as a positive control for leaf senescence. When the detached rosette leaves from 3-week-old *Arabidopsis* were treated in the dark for 4 days, the leaf phenotypes of *ATG6-GFP* and *ATG6-mCherry* were greener than Col and the chlorophyll content in *ATG6-GFP* and *ATG6-mCherry* was also significantly higher than Col (**Fig. S7**). The severity of senescence followed the order: *atg5* > amiRNA*^ATG6^* # 2 > amiRNA*^ATG6^* # 1 > Col > *ATG6-GFP* or *ATG6-mCherry* (**Fig. S7**). These results suggest that overexpression of *ATG6* delays carbon starvation-induced leaf senescence, and ATG6-GFP and ATG6-mCherry fusion proteins are functional.

**Supplemental Methods**

**Methods S1. Plasmid construction**

*For plant transformation*

The *ATG6* coding regions were prepared by PCR with Ex Taq DNA polymerase (TaKaRa, RR001A, Dalian, China) and cloned into 1300-UBQ-mCherry or 1300-UBQ-GFP via double digests and T4 DNA ligase. The amiRNA*^ATG6^* recombinant plasmid was constructed by double digestion. The amplification primers for the amiRNA*^ATG6^* precursor, including miR-s (primer I), miR-a (primer II), miR*s (primer III), and miR*a (primer IV), were designed using the artificial microRNA (amiRNA) design platform Web MicroRNA Designer (WMD3, http://wmd3.weigelworld.org). To generate the stem-loop structure of the amiRNA*^ATG6^* precursor, pCB302-amiR-GFP was utilized as a template (Zhang *et al.*, 2018). An overlap extension PCR method (Niu *et al.*, 2006; Carbonell *et al.*, 2015), was employed for the synthesis of amiRNA*^ATG6^*. The specific procedure involved two rounds of amplification. Firstly, miR319 F and primer IV, miR319 R and primer I, primer II and primer III were used for the first round of amplification. Subsequently, a combination of the first-round PCR amplification products served as templates for the second round of PCR amplification using miR319 F and miR319 R primers. The resulting products from the second-round PCR were digested with appropriate restriction endonucleases (BamH I and Sma I) and then ligated to the 1300-UBQ-mCherry and pERM10M vectors using T4 DNA ligase.

*For Yeast two-hybrid assay*

For the interaction of ATG6 and NPR1.The coding regions of *ATG6*, *SnRK2.8*, *NPR1*, *NPR1-N* (1~984 bp), *NPR1-C* (984~1782 bp) were prepared by PCR with Ex Taq DNA polymerase (TaKaRa, RR001A, Dalian, China) using the primers containing 15~20 bp homologous sequence of the linearized pGADT7 or pGBKT7 vector. The *ATG6* and *SnRK2.8* coding regions were cloned into pGADT7 via ClonExpress II One Step Cloning Kit (Vazyme, C112-02, Nanjing, China); the coding regions of *NPR1*, *NPR1-N* and *NPR1-C* were cloned into pGBKT7, respectively. For interaction of ATGs and NPRs. The coding region of ATGs and NPRs was generated by PCR with Ex Taq DNA polymerase using the primers containing Gateway attB sites. The amplified fragment was cloned into the pDONR207 vector by the BP Clonase II reaction (Invitrogen, 11789-020, Waltham, MA, USA). Each positive clone was inserted into the gateway destination pDEST-GBKT7 and pDEST-GADT7 for yeast transformation by LR Clonase II (Invitrogen, 11791-020, Waltham, MA, USA).

*For pull down assay*

The *ATG6* and *SnRK2.8* coding regions were generated by PCR with Ex Taq DNA polymerase (TaKaRa, RR001A, Dalian, China) and cloned into pGEX-4T-1 via double digests (pGEX-4T-1 was provided by Dr. Sheng Li of South China Normal University) and T4 DNA ligase; NPR1-His (pET32a-NPR1) was provided by Dr. ZhengQing Fu of University of South Carolina.

*For the bimolecular fluorescence complementation assay*

The coding regions of *ATG6*, *SnRK2.8* and *NPR1* were generated by PCR with Ex Taq DNA polymerase (TaKaRa, RR001A, Dalian, China) using the primers containing a homologous sequence (15~20 bp) of linearized 35s-gene-nYFP or 35s-gene-cYFP vector. The *ATG6* and *SnRK2.8* coding regions were cloned into 35s-gene-cYFP via the ClonExpress II One Step Cloning Kit (Vazyme, C112-02, Nanjing, China); NPR1 were cloned into 35s-gene-nYFP.

**Methods S2. *Arabidopsis* *thaliana* screening.**

For *UBQ10::ATG6-mCherry* and *UBQ10::*ATG6-GFP plants, *Agrobacterium tumefaciens* strain GV3101 harboring ATG6-mCherry and ATG6-GFP was used for Col transformation. *Agrobacterium tumefaciens* strain GV3101 harboring ATG6-mCherry and ATG6-GFP was used for Col transformation. *Agrobacterium tumefaciens* was cultured in LB solid medium containing 25 mg/L rifampicin (rif) and 50 mg/L kanamycin (kana) for 2 days, and then a single clone was grown in LB liquid medium containing 25 mg/L rif and 50 mg/L kana for 16~18 h at 28℃, 180 rpm. Centrifuge the bacteria at 4000 g for 10 mins, then resuspend the bacteria in 100 mL of permeate (5% sucrose, 0.05% Silwet-77, mix well before dipping, OD_600_ = 0.8~1.0). The mossy flowering plants were selected and the pods and pollinated flowers were removed before transformation. The inflorescence of *Arabidopsis* was placed in the transformation medium containing *Agrobacterium tumefaciens* for 1 min. Incubate and moisturize in the dark for 2 days. Cultivation was continued until plants matured and seeds were collected to screen positive plants. Positive plants with 30 mg/L hygromycin B to homozygous T_3_ lines.

For the generation of amiRNA*^ATG6^* Lines, we utilized *Agrobacterium tumefaciens* strain GV3101 containing the stem-loop structure of the amiRNA*^ATG6^* precursor for the transformation of Col plants. The transformation procedure employed was the same as that used for *ATG6-GFP* plants. Positive plants with 50 mg/L kana to homozygous T_3_ lines.

*UBQ10::ATG6-mCherry* × *35S::NPR1-GFP/npr1-2* (*ATG6-mCherry* x *NPR1-GFP*) was obtained by crossing female *NPR1-GFP/npr1-2* with *ATG6-mCherry*. Screen positive plants with 50 mg/L kana and 30 mg/L hygromycin B. To visualize the localization of NPR1 and ATG6, positive plants with GFP (excitation at 488 nm wavelengths, detection of 500~550 nm wavelengths) and mCherry (excitation at 561 nm wavelengths, detection of 570~650 nm wavelengths) fluorescence were screened through laser scanning confocal microscopy (Zeiss LSM880). To avoid the possibility that the observed fluorescence was due to free mCherry and free GFP, we also verified the presence of ATG6-mCherry and NPR1-GFP in *ATG6-mCherry* x *NPR1-GFP* plants using western blot experiments. In addition, the levels of NPR1-GFP and free GFP in *ATG6-mCherry* x *NPR1-GFP* plants were detected before and after SA treatment. Only ~ 10 % of free GFP was detected in *ATG6-mCherry* x *NPR1-GFP* plants before and after SA treatment (Fig S4). This also means that the fluorescence signal observed by laser scanning confocal microscopy is dominated by NPR1-GFP, not free GFP. For the identification of *npr1-2*, PCR was performed according to the following primer pairs F: GGATGATTTCTACAGCGACGCT, R: GTAACCATAGCTTA ATGCAGATGGTG. PCR procedure as follows. 95℃ 5 min, 95℃ 30 s, 55℃ 30 s, 72℃ 30 s, 72℃ 5 min, 30 cycles. The PCR product was then digested by FspI (R0135V, NEB) at 37℃ for 10 min (reaction system: 20 µL PCR produces, 3 µL NEB Buffer, 0.3 µL FspI and 6.7µL H_2_O) and analyzed by 3% agarose gel electrophoresis (Cao *et al.*, 1997; Chen *et al.*, 2021a), by the same method plants were screened to T_3_.

*NPR1-GFP*/*atg5* was obtained by crossing. For the identification of *atg5*, the triple primer PCR method was performed using RP+LP and LB+RP according to the following primer pairs, LP, AAAGACCACAGAACCCGAAAC. RP, CCAAATTGAATCTTCACCAGG. LBb1.3, ATTTTGCCGATTTCGGAAC. Screen positive plants with 50 mg/L kana, and then positive plants with GFP fluorescence were screened through laser scanning confocal microscopy. Then, in order to exclude the possible fluorescence effects of free GFP, we also used western blots to verify the presence of NPR1-GFP until to T_3_ homozygous lines.

**Methods S3. For *Pst* DC3000/*avrRps4* culture and infiltration.**

*Pst* DC3000/*avrRps4* was grown in KB medium containing 50 mg/L kana and 25 mg/L rif for 18~24 h at 28℃, 180 rpm. Then centrifuged at 4000 g for 10 mins. The precipitate was washed twice with 10 mM magnesium chloride (MgCl_2_) and resuspended. The absorbance of the suspension liquid was measured at 600 nm and gradually diluted from 0.8 to 0.02 (for protein levels) or 0.001 (for growth of pathogenic bacteria). Infiltration with *Pst* DC3000/*avrRps4* was performed by pressure infiltration with a 1 mL syringe through the abaxial leaf surface.

**Methods S4. Yeast two-hybrid assay.**

For the interaction of ATG6 with NPR1, the plasmids combinations NPR1-BD and ATG6-AD, NPR1-N-BD and ATG6-AD, NPR1-C-BD and ATG6-AD were respectively co-transformed into the yeast strain AH109, according to the Clontech yeast transformation protocol. Co-transformation plasmid combinations NPR1-BD and AD, BD and ATG6-AD, NPR1-N-BD and AD, NPR1-C-BD, SnRK2.8-AD and BD and AD as negative controls. Co-transformation of NPR1-BD and SnRK2.8-AD used as positive control. Yeast strains were cultivated on SD/-Trp-Leu for 3 days. Pick a fresh single clone and add 50 μL of SD-2 liquid media. Then, yeast strains are gradually diluted in 3 gradients (10^-1^, 10^-2^, 10^-3^). The yeast was added to a SD/-Trp-Leu-His-Ade to analyze the interaction.

For the interaction of ATGs and NPRs, pGBKT7-NPRs (NPRs-BD) and pGADT7-ATGs (ATGs-AD) were co-transformed into the yeast strain AH109. Yeast strains were cultured on SD/-Trp-Leu for 3 days. Pick a fresh single clone and add it to 50 μL of SD-2 liquid media. Then yeast was added to a SD/-Trp-Leu-His-Ade with different SA concentrations to analyze their interaction.

**Methods S5.** **Prokaryotic protein expression.**

*E. coli strain* BL21 (DE3) harboring GST, GST-ATG6, GST-SnRK2.8 and NPR1-His was cultured in LB solid medium with 50 mg/L Ampicillin (Amp) for 12~16 h. Single clones were selected and grown in 1 mL liquid LB medium overnight at 37℃, 150 rpm. Transfer 1 mL of the bacterial solution to 100 mL of LB medium and incubate until OD_600_ = 0.6~1.0. NPR1-His bacterial solution was induced by 1 mM Isopropyl β-D-Thiogalactoside (IPTG) at 16°C for 24 h, 150 rpm; GST, GST-ATG6 and GST-SnRK2.8 bacterial solution were induced by 1 mM IPTG at 16°C for 6 h, 150 rpm. Then it is centrifuged at 8422 g for 10 mins at 4°C. The NPR1-His precipitate was added to 4~5 times the volume of Ni-buffer A (20 mM Tris-HCl pH 7.5, 300 mM NaCl, 15 mM imidazole, 1 mM β-Mercaptoethanol). The precipitate of GST, GST-ATG6, GST-SnRK2.8 was added to 4~5 times the volume of PBS buffer (10 mM Na_2_HPO_4_, 140 mM NaCl, 2.7 mM KCl, 1.8 mM KH_2_PO_4_, pH 7.4), and added to the cells were dissolved by ultrasound. Centrifuge at 8422 g for 20 mins and aspirate the supernatant.

**Methods S6. Nuclear and cytoplasmic separation of NPR1-GFP.**

Samples (0.5 g) of *Arabidopsis* or *Nicotiana benthamiana* were ground well into powder in liquid nitrogen and then added to 1 mL Hondar buffer (2.5% Ficoll 400, 5% DextranT 400, 0.4 M sucrose, 25 mM Tris-HCl pH 7.5, 10 mM MgCl_2_, 10 mM β-Mercaptoethanol) containing freshly configured 0.5 mM PMSF (329-98-6, Sigma-Aldrich, USA), 40 μM MG115 (47480, Sigma-Aldrich, USA), 500 × protease inhibitor cocktail (Aprotinin 500 μg/mL, Leupeptin 500 μg/mL, Pepstatin 500 μg/mL) and 5000 × phosphatase inhibitor cocktail (Na_3_VO_4_·12H_2_O 10 μg/mL, NaF 2.5 μg/mL). The tissue solution was filtered through a 62 μm nylon mesh filter and then a final concentration of 0.5% Triton-100 was added to the filtrate. The mixture was gently mixed and incubated on ice for 15 mins. Centrifuged at 1500 g for 5 mins at 4°C. The supernatant (cytoplasmic fraction) was carefully transferred to a new centrifuge tube, while the precipitate was resuspended in 1 mL Hondar buffer containing 1% Triton-100. Centrifugated at 100 g for 1 min at 4°C to remove residual cytoplasmic fractions. Then the upper layer was aspirated and centrifuged at 1500 g for 5 mins to obtain the nuclear fraction. The nuclear fraction was washed three times with Hondar buffer, and 100 μL of Hondar buffer was added to dissolve it. The protein concentration of the samples was determined using a Bradford microplate reader (INFINITE M PLEX, Tecan). After 10 mins denaturation at 75°C, Western blot was performed.

**Methods S7. Protein Degradation Analysis.**

Samples (0.5 g) of *Arabidopsis* (*NPR1-GFP* and *ATG6-mCherry × NPR1-GFP*) or *Nicotiana benthamiana* were fully ground in liquid nitrogen and mixed with 500 μL of basal buffer without protease inhibitors (100 mM Tris-HCl pH 7.5, 1 mM EDTA, 150 mM NaCl, 0.5% (v/v) Nonidet P-40). The samples were divided equally into 6 portions and treated at 25°C for 0, 30, 60, 120 and 180 mins (Spoel *et al.*, 2009). A sample was treated with 100 μM MG115 for 180 mins to inhibit the proteasome degradation pathway. One of the portions was treated with 5 μM concanamycin A (Invitrogen, Waltham, MA, USA) or 30 μM Wortmannin (19545–26–7, MedChemExpress, NJ, USA) for 120 mins to inhibit autophagy. The protein samples were denatured at 75°C for 10 mins. Subsequently, the samples were subjected to SDS-polyacrylamide gel electrophoresis (SDS-PAGE) electrophoresis and analyzed according to Western blotting.

**Methods S8. Protein Extraction and Western Blotting.**

Leaves or seedlings (400 mg) were fully ground in liquid nitrogen and homogenized in 400 μL basal buffer (100 mM Tris-HCl pH 7.5, 1 mM EDTA, 150 mM NaCl, 0.5% (v/v) Nonidet P-40, 1 mM PMSF). Add 40 μM MG115, 500 × protease inhibitor cocktail and 5000 × phosphatase inhibitor cocktail to the basal buffer. The samples were incubated on ice for 30 mins, with vortexed every 10 mins, followed by centrifuged at 10142 g (TGL16, cence, hunan, China) for 15 mins at 4°C. The supernatant was transferred to a new 1.5 mL centrifuge tube. The protein concentration of the samples was determined by Bradford using a microplate reader (INFINITE M PLEX, Tecan). Protein was denatured at 100°C for 10 mins. NPR1 protein was denatured at 75°C for 10 mins.

The sample were subjected to SDS-PAGE (10%) and the gel was transferred to 0.45 μM (> 35 KDa) or 0.22 μM (<35 KDa) PVDF membrane (IPVH00010, Merck Millipore, Germany) for 60 mins (> 35 KDa) or 30 mins (<35 KDa). The membrane was blocked in TBST (Tris-buffered saline with Tween-20) containing 5% dry milk at room temperature for 2 h. After three washes with TBST, the membrane was incubated with the primary antibody overnight at 4°C. Then, the membranes were washed three times with TBST, and incubated with secondary antibody at room temperature for 2 h. Finally, the membrane was washed three times with TBST and the chemiluminescence was imaged using an image analyzer (Tanon-5200, Shanghai, China).

**Methods S9. For the treatment of 3-week-old *Nicotiana benthamiana*.**

*Agrobacterium* was initially cultured in LB solid medium (contains appropriate antibiotics). A single clone was then selected and grown in 30 mL of LB liquid medium with the same antibiotics for 16~18 h at 28℃, 180 rpm. The supernatant was removed by centrifugation and the precipitate was suspended in the infiltration buffer (10 mM MES pH 5.7, 10 mM MgCl_2_, 150 µM Acetosyringone). The absorbance of the suspension liquid at 600 nm was gradually diluted to OD_600_ = 0.5~0.8. The diluted suspension was mixed in a 1:1 ratio. After incubation at 25℃ for 1 h, the mixed *Agrobacterium* was infiltrated into leaves of 3-week-old *Nicotiana benthamiana* using a previously described method (Jiao *et al.*, 2019).

**Methods S****10. For the bimolecular fluorescence complementation assay.**

The *Agrobacterium* GV3101 mixture, containing various combinations of ATG6-cYFP and NPR1-nYFP, NPR1-nYFP and SnRK2.8-cYFP, nYFP and ATG6-cYFP, NPR1-nYFP and cYFP, and nYFP and SnRK2.8-cYFP, was infiltrated into nls-mCherry transgenic tobacco. After 3 days of infiltration, YFP fluorescence was detected in epidermal cells by laser scanning confocal microscopy using an excitation wavelength of 518 nm, detection at 500~550 nm wavelengths. mCherry was detected using an excitation of 561 nm, detection of 570~650 nm wavelengths.

**Methods S11. For** **SINCs-like condensates observation.**

*Agrobacterium* strain GV3101 harboring NPR1-GFP and mCherry; NPR1-GFP and ATG6-mCherry were mixed in a 1:1 ratio. After incubation at 25℃ for 1 h, the mixed *Agrobacterium* was infiltrated into leaves of 3-week-old *Nicotiana benthamiana*. After 2 d, the leaves were soaked in 1 mM SA solution for 24 h. And then GFP (excitation of 488 nm wavelengths, detection of 500-550 nm wavelengths) fluorescence signals were observed under the laser scanning confocal microscopy.

**Methods S12. For growth of *Pst* DC3000*/avrRps4***

A low dose (OD_600_ = 0.001) of *Pst* DC3000/*avrRps4* was infiltrated. After 3 days, two small round leaves (8 mm in diameter) were ground into powder in a 1.5 mL centrifuge tube, and then added 500 μL of MgCl_2_ to dissolve it as the original solution. The original liquid is gradually diluted in 6 gradients (10^-1^, 10^-2^, 10^-3^, 10^-4^, 10^-5^, 10^-6^). *Pst* DC3000/*avrRps4* were spread onto KB solid media (containing 25 mg/L rif and 50 mg/L kana). After 2 days, the colony number was counted according to a previous description (Wang *et al.*, 2016; Lei *et al.*, 2020).

**Methods S13. Real-Time Quantitative PCR (RT-qPCR).**

Total RNA was extracted using Trizol RNA reagent (Invitrogen, 10296–028, Waltham, MA, USA). cDNA was synthesized from 1 µg high-quality total RNA using RT Reagent Kit (TaKaRa, RR047A, Dalian, China). The qPCR was performed on ABI Life QuantStudio 6 using the Low ROX Premixed ChamQ SYBR qPCR Master Mix (Vazyme, Q331-02, Nanjing, China). The qPCR thermal cycles were as follows: 95°C for 30 s, followed by 40 cycles of 95°C for 5 s and 60 °C for 34 s. *AtActin2* was used as a control. Each reaction was independently repeated at least three times. Relative expression levels of target genes were calculated using the relative 2^−∆∆Ct^ method (Zhang *et al.*, 2023).

**Accession numbers**

Sequence data in this article can be found in the GenBank/TAIR databases under the following accession numbers: *Actin2*, At3G18780; *NPR1*, AT1G64280; *ATG6*, AT3G61710; *SnRK2.8*, AT1G78290; *PR1*, AT2G14610; *PR5*, AT1G75040; *ICS1*, AT1G74710.

**Supplemental Moves**

**Supplemental Move 1.** Localization of NPR1-GFP in *Nicotiana benthamiana* leaves co-expressed NPR1-GFP and mCherry.

**Supplemental Move 2.** Localization of NPR1-GFP in *Nicotiana benthamiana* leaves co-expressed NPR1-GFP and ATG6-mCherry.
