## Supplemental Data 2 for "ATG6 interacting with NPR1 increases *Arabidopsis thaliana* resistance to *Pst* DC3000/*avrRps4* by increasing its nuclear accumulation and stability"

**The following supporting information is available for this article**

**Vector mapping used in this study.**

Table 1. Vector mapping used in this study

| Number | Name | Hyperlink |
| --- | --- | --- |
| 1 | pCB302-GFP | Fig. 1 |
| 2 | pGADT7 | Fig. 2 |
| 3 | pGBKT7 | Fig. 3 |
| 4 | pGEX-4T-1 | Fig. 4 |
| 5 | pET-32a (+) | Fig. 5 |
| 6 | pCAMBIA1300 UBQ-sGFP | Fig. 6 |
| 7 | pCAMBIA1300 UBQ-mCherry | Fig. 7 |
| 8 | YC-BIFC-gene-cYFP | Fig. 8 |
| 9  10 | YC-BIFC-gene-nYFP  pERM10 | Fig. 9  Fig. 10 |

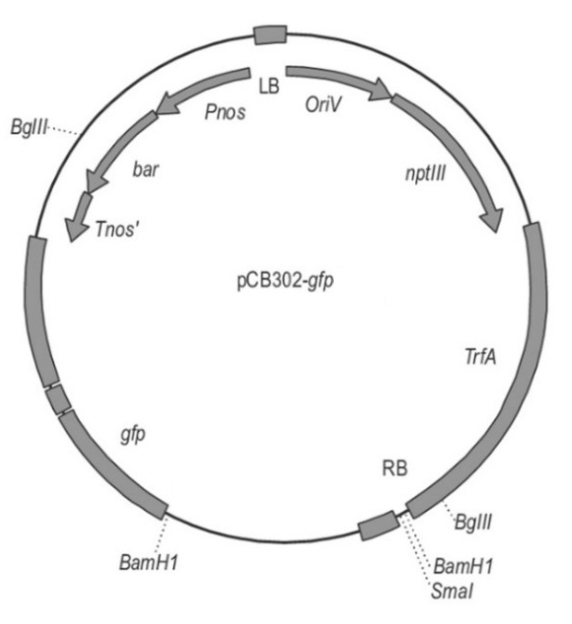

Supplemental Data 2 Figure 1. Map of pCB302-GFP vector.

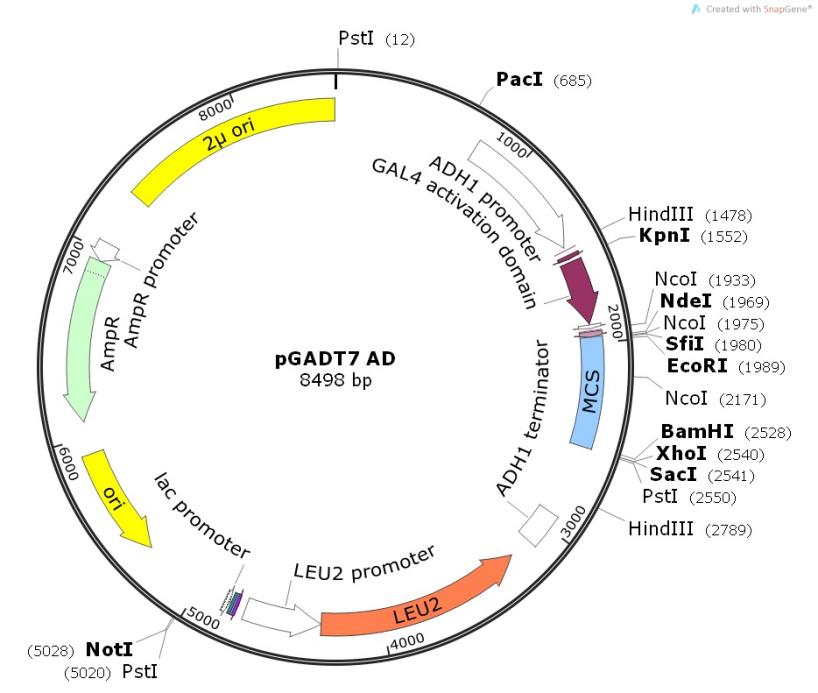

Supplemental Data 2 Figure 2. Map of pGADT7 AD vector.

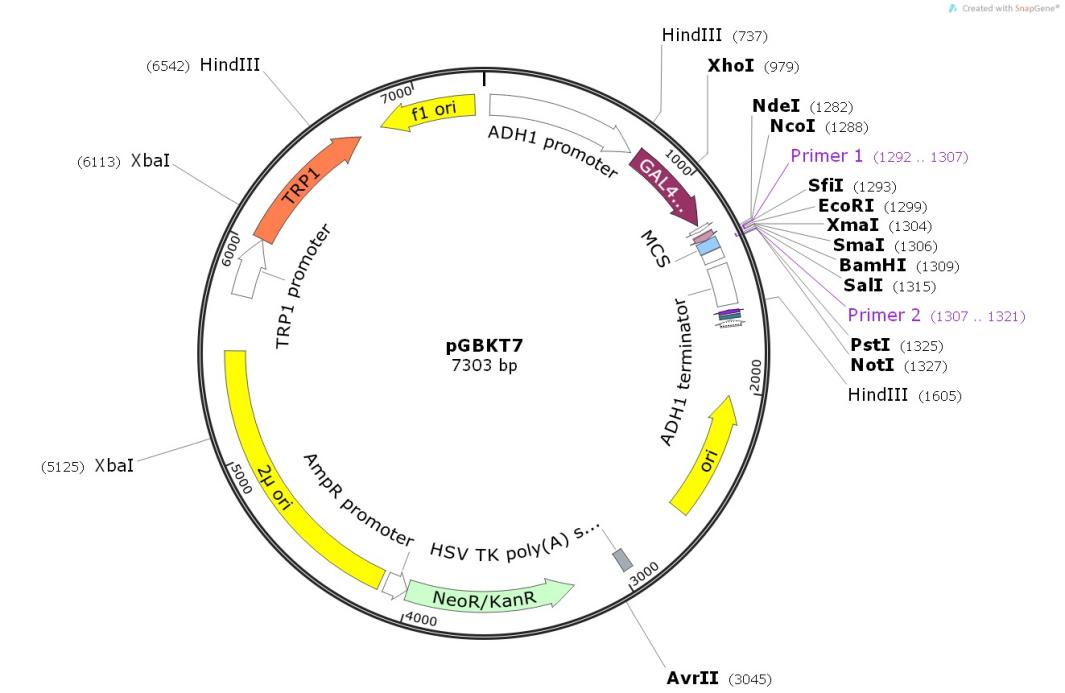

Supplemental Data 2 Figure 3. Map of pGBKT7 BD vector.

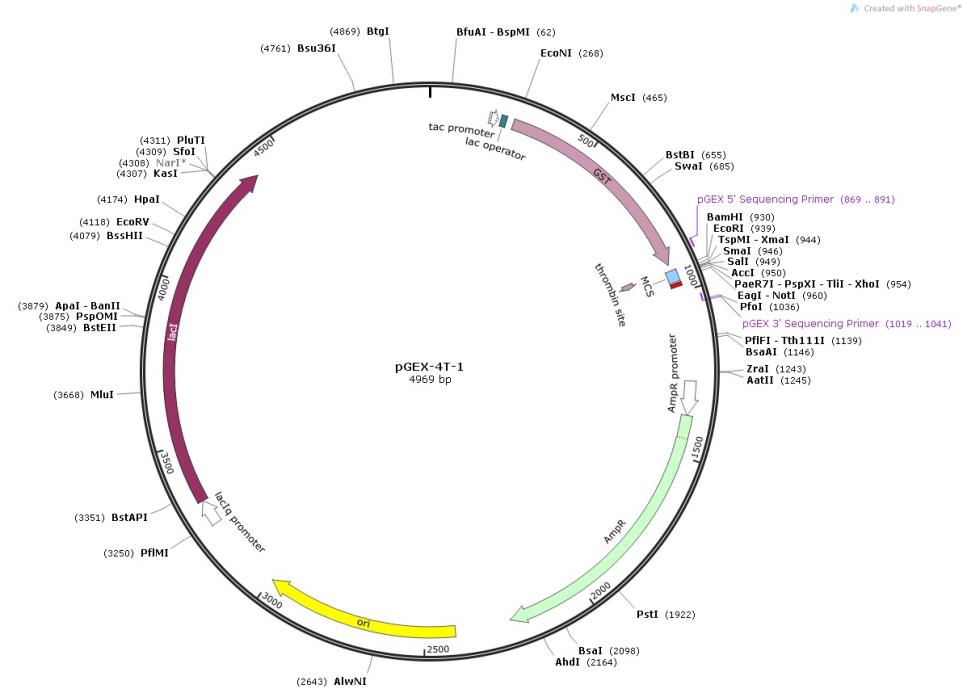

Supplemental Data 2 Figure 4. Map of pGEX-4T-1 vector.

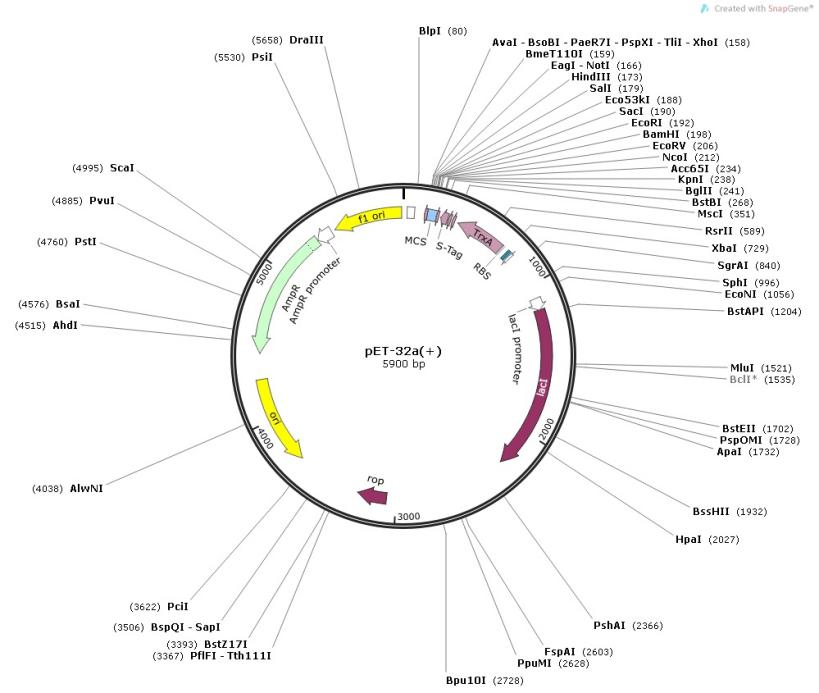

Supplemental Data 2 Figure 5. Map of pET-32a(+) vector.

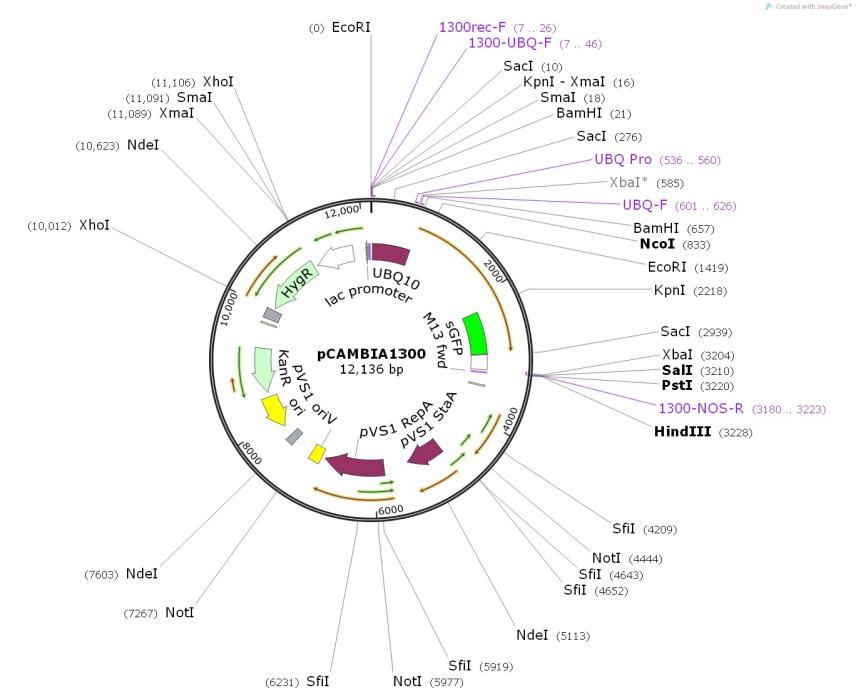

Supplemental Data 2 Figure 6. Map of pCAMBIA1300 UBQ-sGFP vector.

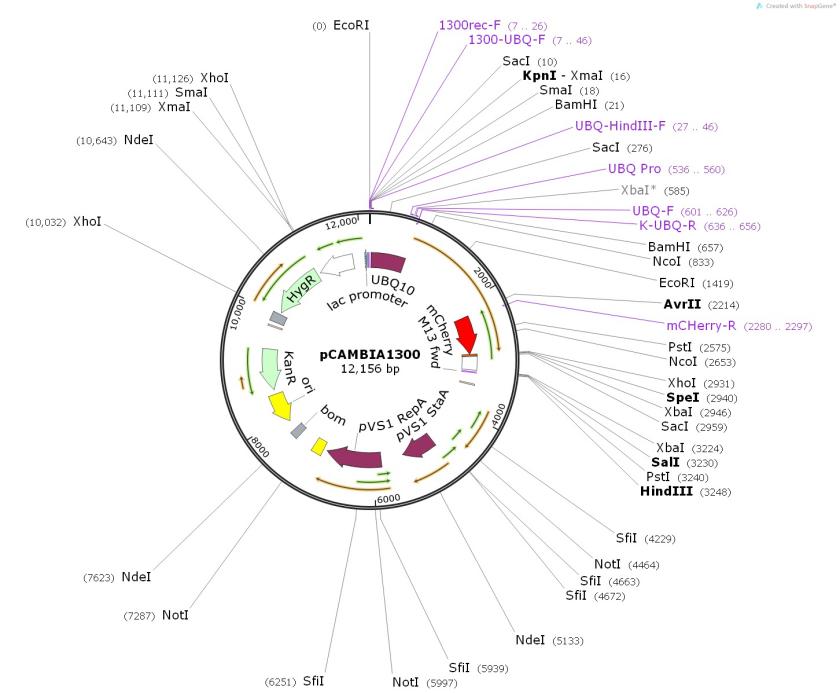

Supplemental Data 2 Figure 7. Map of pCAMBIA1300 UBQ-mCherry vector.

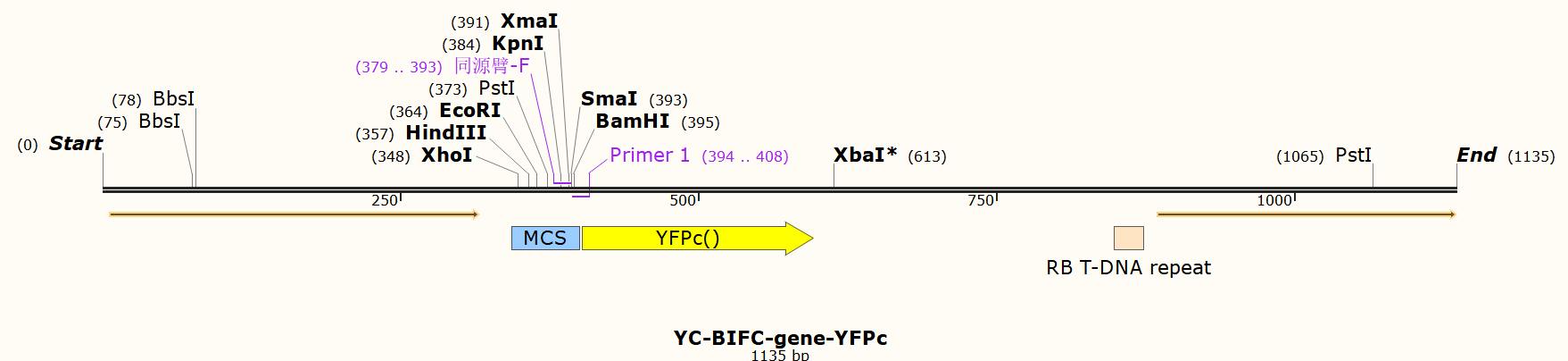

Supplemental Data 2 Figure 8. Map of YC-BIFC-gene-cYFP vector.

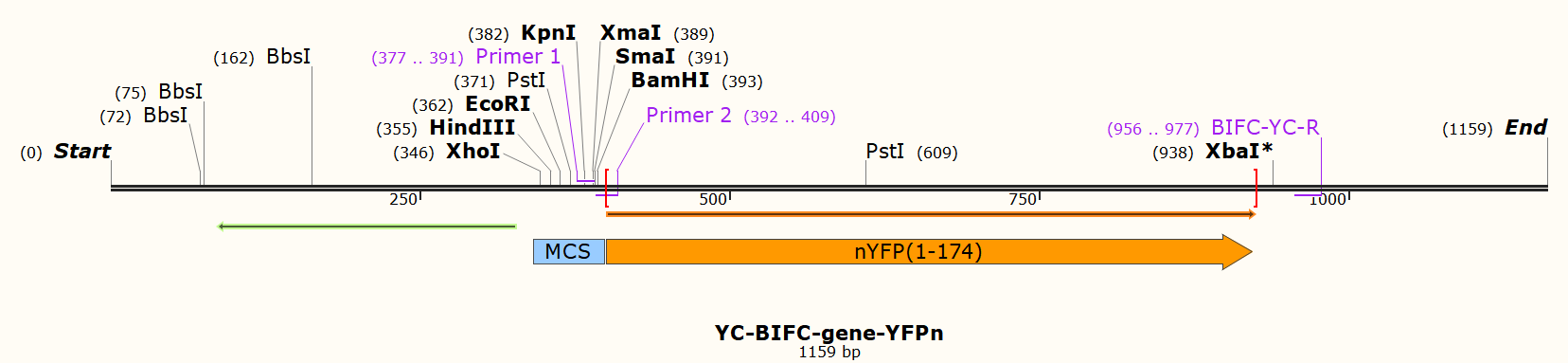

Supplemental Data 2 Figure 9. Map of YC-BIFC-gene-nYFP vector.

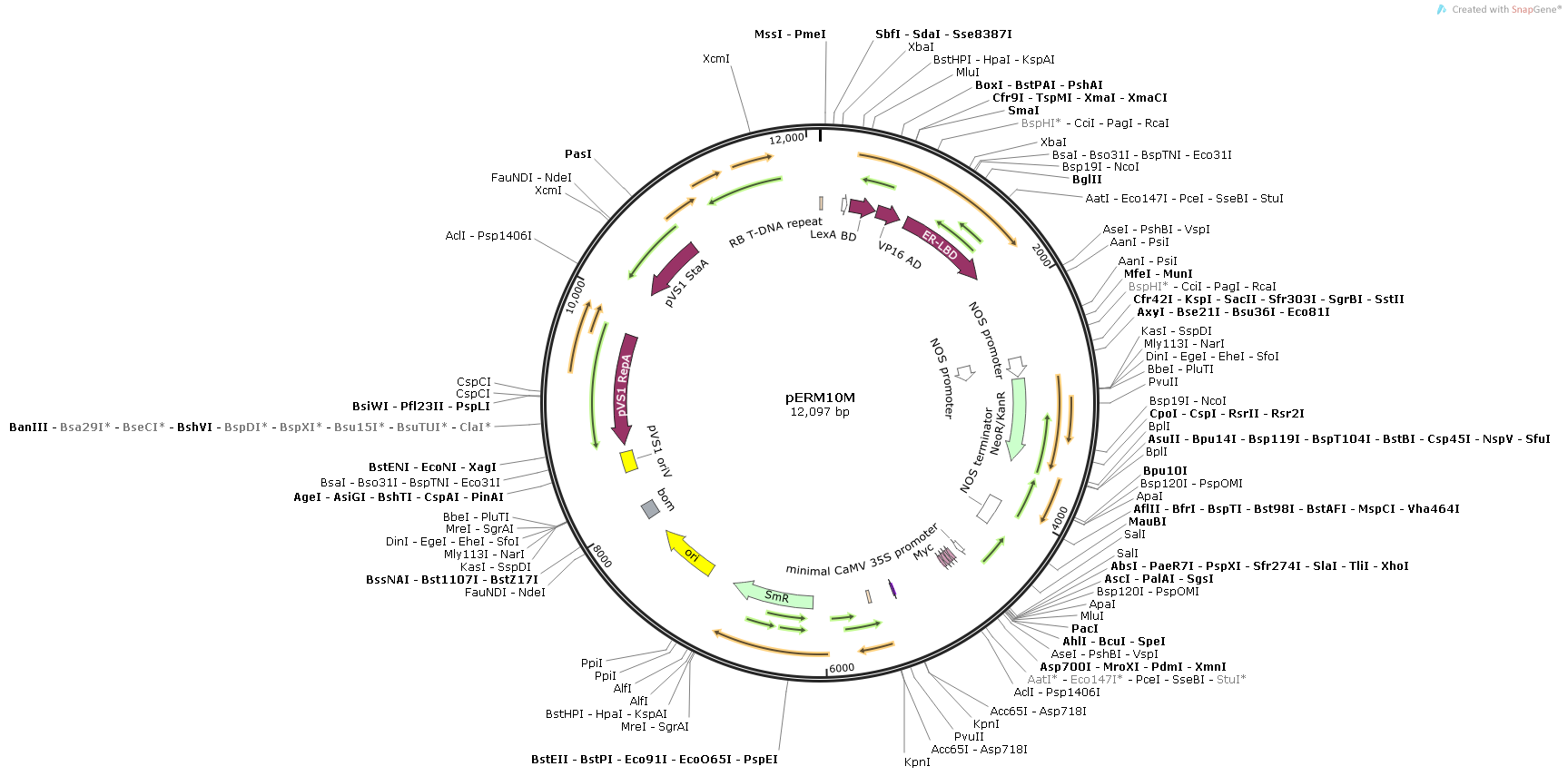

Supplemental Data 2 Figure 10. Map of pERM10 vector.
